# sumrep: a summary statistic framework for immune receptor repertoire comparison and model validation

**DOI:** 10.1101/727784

**Authors:** Branden J Olson, Pejvak Moghimi, Chaim Schramm, Anna Obraztsova, Duncan Ralph, Jason A Vander Heiden, Mikhail Shugay, Adrian Shepherd, William Lees, Frederick A Matsen

## Abstract

The adaptive immune system generates an incredible diversity of antigen receptors for B and T cells to keep dangerous pathogens at bay. The DNA sequences coding for these receptors arise by a complex recombination process followed by a series of productivity-based filters, as well as affinity maturation for B cells, giving considerable diversity to the circulating pool of receptor sequences. Although these datasets hold considerable promise for medical and public health applications, the complex structure of the resulting adaptive immune receptor repertoire sequencing (AIRR-seq) datasets makes analysis difficult. In this paper we introduce `sumrep`, an R package that efficiently performs a wide variety of repertoire summaries and comparisons, and show how `sumrep` can be used to perform model validation. We find that summaries vary in their ability to differentiate between datasets, although many are able to distinguish between covariates such as donor, timepoint, and cell type for BCR and TCR repertoires. We show that deletion and insertion lengths resulting from V(D)J recombination tend to be more discriminative characterizations of a repertoire than summaries that describe the amino acid composition of the CDR3 region. We also find that state-of-the-art generative models excel at recapitulating gene usage and recombination statistics in a given experimental repertoire, but struggle to capture many physiochemical properties of real repertoires.

## Introduction

B cells and T cells play critical roles in adaptive immunity through the cooperative identification of, and response to, antigens. The random rearrangement process of the genes that construct B cell receptors (BCRs) and T cell receptors (TCRs) allows for the recognition of a highly diverse set of antigen epitopes. We refer to the set of B and T cell receptors present in an individual’s immune system as their immune receptor repertoire; this dynamic repertoire constantly changes over the course of an individual’s lifetime due to antigen exposure and the effects of aging.

Although immune receptor repertoires are now accessible for scientific research and medical applications through high-throughput sequencing, it is not necessarily straightforward to gain insight from and to compare these datasets. Indeed, if these datasets are not processed, they are simply a list of DNA sequences. After annotation one can compare gene usage [15, 21, 7, 11, 5, 4] and CDR3 sequences. This can be a highly involved task, and so it is common to simply compare the gene usage frequencies and CDR3 length distributions of repertoire [23, 17], leaving the full richness of the CDR3 sequence and potentially interesting aspects of the germline-encoded regions unanalyzed.

An alternative strategy is to transform a repertoire to a more convenient space and compare the transformed quantities according to some metric. For example, several studies reduce a set of nucleotide sequences to *k*mer distributions for classification of immunization status or disease exposure [19, 26, 14]. These *k*mer distributions can then be compared via a string metric, but still comprise a large space and lose important positional information. One can perform other dimension reduction techniques such as t-SNE to project repertoires down to an even smaller space [39], but these projections also discard a lot of information and can be difficult to interpret biologically.

We wish to facilitate the use of biologically interpretable summary statistics to capture many different aspects of AIRR-seq data. In addition to enabling comparison of different sequencing datasets, summary statistics can also be used to compare sequencing datasets to probabilistic models to which they have been fitted. Namely, one can use a form of model checking that is common in statistics: after fitting a model to data, one assesses the similarity of the model-generated data to the real data. In this case, we generate a repertoire of sequences from models and compare this collection to a real-data repertoire of sequences via summary statistics.

We are motivated to perform such comparison because these probabilistic models are used as part of inference, and because they are used for inferential tool benchmarking. Such generative models are used to simulate sequences as a “ground truth” for benchmarking inferential software [31, 12, 20], and thus the accuracy of such benchmarks depends on the realism of the generated sequences. Simulation tools can also be used to generate a null distribution used to test for a specific effect, such as natural selection [37].

Currently, there are no unified packages dedicated to the task of calculating and comparing summary statistics for AIRR-seq datasets. While the Immcantation framework (which includes the `shazam` and `alakazam` R packages) contains many summary functions for AIRR-seq data [13], it does not have general functionality for retrieving, comparing, and plotting these summaries. Many summaries of interest are implemented in one package or another, but differences in functionality and data structures make it troublesome to compute and compare summaries across packages. Some summaries of interest, such as the distribution of positional distances between mutations, are not readily implemented in any package.

In this paper, we gather dozens of meaningful summary statistics on repertoires, derive efficient and robust summary implementations, and identify appropriate comparison methods for each summary. We present `sumrep`, an R package that computes these summary distributions for AIRR-seq datasets and performs repertoire comparisons based on these summaries. We investigate the effectiveness of various summary statistics in distinguishing between different experimental repertoires as well as between simulated and experimental data. We show that many summaries differentiate between various covariates by which the datasets are stratified. Further, we demonstrate how `sumrep` can be used for model validation through case studies of two state-of-the-art repertoire simulation tools: `IGoR` [20] applied to TRB sequences, and `partis` [30, 31] applied to IGH sequences.

## Results

### Implementation

The full `sumrep` package along with the following analyses can be found at https://github.com/matsengrp/sumrep. It supports the IGH, IGK, and IGL loci for BCR datasets, and the TRA, TRB, TRD, and TRG loci for TCR datasets. It is open-source, unit-tested, and extensively documented, and uses default dataset fields and definitions that comply with the Adaptive Immune Receptor Repertoire (AIRR) Community Rearrangement schema [36]. A reproducible installation procedure of `sumrep` is available using Docker [3].

Table 1 lists the summary statistics currently supported by `sumrep`, and includes the default assumed degree of annotation, clustering, and phylogenetic inference for each summary. The first group of statistics only requires the input or query sequences to be aligned to their inferred germline sequences (e.g. IMGT-aligned) and constrained to the variable region; this coincides with the presence of the `sequence_alignment` and `germline_alignment` fields in the AIRR schema. The second group requires standard sequence annotations, such as inferred germline ancestor sequences for Ig loci, germline gene assignments, and indel statistics. The third group requires clonal family cluster assignments. The fourth group requires a inferred phylogeny for each clonal family of an Ig dataset. `sumrep` itself does not perform any annotation, clustering, or phylogenetic inference, but rather assumes such metadata are present in the given dataset; in principle, one can use any tool which performs these tasks as expected.

**Table 1:** Currently supported summary statistics grouped by their respective degrees of assumed post-processing. Annotation denotes whether annotation of the V(D)J germline segment is required, Clustering denotes whether clonal clustering is required, and Phylogeny denotes whether lineage tree inference is required.

| Summary statistic | Annotations | Clustering | Phylogeny | Implementation |
| --- | --- | --- | --- | --- |
| Pairwise distance distribution | No | No | No | `stringdist` [35] |
| $k$th nearest neighbor distribution | No | No | No | `stringdist` |
| GC-content distribution | No | No | No | `ape` [28] |
| Hotspot motif count distribution | No | No | No | `Biostrings` [27] |
| Coldspot motif count distribution | No | No | No | `Biostrings` [27] |
| CDR3 length distribution | Yes | No | No | Tool-provided |
| Joint distribution of germline gene use | Yes | No | No | `sumrep` |
| Pairwise CDR3 distance distribution | Yes | No | No | `stringdist` |
| Atchley factor distributions | Yes | No | No | `HDMD` [22] |
| Kidera factor distributions | Yes | No | No | `Peptides` [22] |
| Aliphatic index distribution | Yes | No | No | `Peptides` |
| G.R.A.V.Y. index distribution | Yes | No | No | `alakazam` [13] |
| Polarity distribution | Yes | No | No | `alakazam` |
| Charge distribution | Yes | No | No | `alakazam` |
| Basicity distribution | Yes | No | No | `alakazam` |
| Acidity distribution | Yes | No | No | `alakazam` |
| Aromaticity distribution | Yes | No | No | `alakazam` |
| Bulkiness distribution | Yes | No | No | `alakazam` |
| Per-gene substitution rate | Yes | No | No | Tool-provided + `sumrep` |
| Per-gene-per-position substitution rate | Yes | No | No | Tool-provided + `sumrep` |
| Per-base mutability model | Yes | No | No | `shazam` [13] |
| Per-base substitution model | Yes | No | No | `shazam` |
| Positional distance between mutations distribution | Yes | No | No | `sumrep` |
| Distance from germline to sequence distribution | Yes | No | No | `stringdist` |
| V gene 3' deletion length distribution | Yes | No | No | Tool-provided |
| V gene 5' deletion length distribution | Yes | No | No | Tool-provided |
| D gene 3' deletion length distribution | Yes | No | No | Tool-provided |
| D gene 5' deletion length distribution | Yes | No | No | Tool-provided |
| J gene 3' deletion length distribution | Yes | No | No | Tool-provided |
| J gene 5' deletion length distribution | Yes | No | No | Tool-provided |
| VD (or VJ) insertion length distribution | Yes | No | No | Tool-provided |
| DJ insertion length distribution | Yes | No | No | Tool-provided |
| VD (or VJ) insertion transition matrix | Yes | No | No | `sumrep` |
| DJ insertion transition matrix | Yes | No | No | `sumrep` |
| V/J in-frame percentage | Yes | No | No | Tool-provided + `sumrep` |
| Cluster size distribution | Yes | Yes | No | Custom |
| Hill numbers (diversity indices) | Yes | Yes | No | `alakazam` |
| Selection estimates | Yes | Yes | No | `shazam` |
| Sackin index distribution | Yes | Yes | Yes | `CollessLike` [24] |
| Colless-like index distribution | Yes | Yes | Yes | `CollessLike` |
| Cophenetic index distribution | Yes | Yes | Yes | `CollessLike` |

`sumrep` makes it easy to compare summary statistics between two repertoires by equipping each summary with an appropriate divergence, or measure of dissimilarity, between instances of a summary. For example, the `getCDR3LengthDistribution` function returns a vector of each sequence’s CDR3 length, and the corresponding compareCDR3LengthDistributions function takes two repertoires and returns a numerical summary of the dissimilarity between these two length distributions. The comparison method depends on the summary, which is discussed further in the Methods section. `sumrep` also includes a `compareRepertoires` function which takes two repertoires and returns as many summary comparisons as befit the data.

Figure 1 illustrates the general framework of comparing summary statistics between two repertoires *R*_1_ and *R*_2_. A given summary *s* is applied separately to *R*_1_ and *R*_2_, which for most summaries yields a distribution of values (Figure 1a). These two resultant distributions can be compared using a divergence 𝒟 that is tailored to the nature of *s* (Figure 1b). We use Jenson-Shannon (JS) divergence to compare scalar distributions (e.g. GC content, CDR3 length), which is a symmetrized version of KL-divergence, a weighted average log-ratio of frequencies widely-used in statistics and machine learning. We use the similarly popular ℓ_1_ divergence to compare categorical distributions (e.g. gene call frequencies, amino acid frequencies), which is a sum of absolute differences of counts.

**Figure 1:**
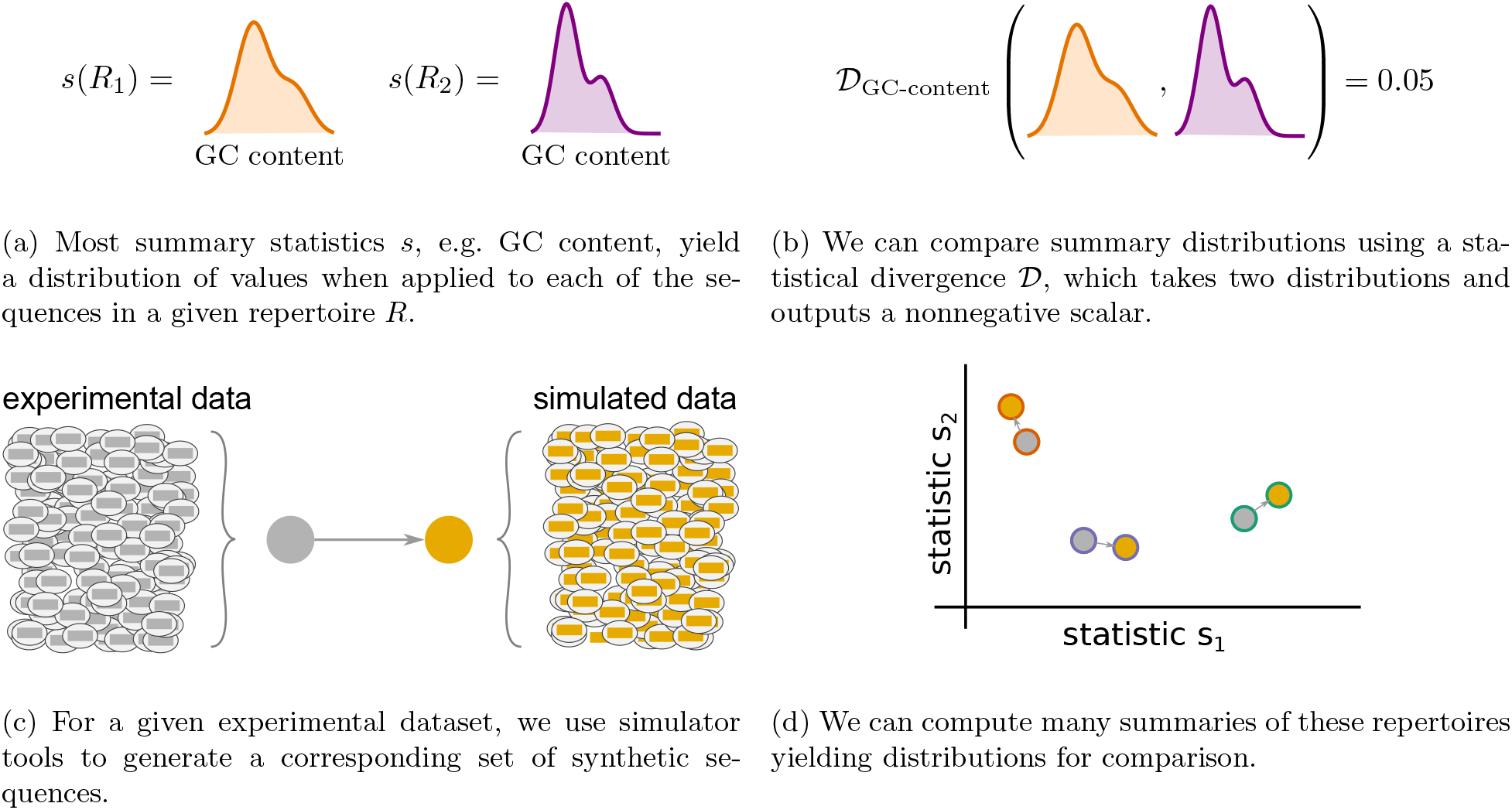
Cartoon of our summary statistic and divergence framework, and how this can be applied to validation of repertoire simulators. Steps (a) and (b) can be applied to compare arbitrary datasets, while (c) and (d) show how sumrep can be used for model comparison.

We have designed `sumrep` to efficiently approximate computationally intensive summaries. When the target summary is a distribution, we can gain efficiency by repeatedly subsampling from the distribution until our estimate has stabilized. The result is an approximation to the full distribution; by introducing slight levels of noise, we can gain very substantial runtime performance improvements for large datasets. This in turn allows fast, accurate divergence estimates between dataset summaries. We outline a generic distribution approximation algorithm as well as a modified version for the nearest neighbor distance distribution in the Methods section, and conduct extensive empirical validation of these algorithms in Appendices A and B.

`sumrep` additionally contains a plotting function for each univariate summary distribution. For example, the `getCDR3LengthDistribution` comes with a companion plotting function called `plotCDR3LengthDistribution`. `sumrep` also includes a master plotting function, `plotUnivariateDistributions`, which shows a gridded figure of all univariate distribution plots relevant to the locus in question which can be computed from the input dataset. Currently, these plotting functions support frequency polygons and empirical cumulative distribution functions (ECDFs). Examples of these plots can be found throughout later sections of this report.

### Application of summary statistics to experimental data

To examine the ability of various summary statistics to distinguish among real repertoires, we applied `sumrep` to TCR and BCR datasets performed a multidimensional scaling (MDS) analysis of summary divergences. In particular, we computed divergences of each summary between each pair of repertoires, stratified by covariates such as individual, timepoint, and cell subset to form a dissimilarity matrix. We then mapped these dissimilarity matrices to an abstract Cartesian space using MDS.

For TCR repertoires, we used datasets from two individuals and five timepoints post-vaccination, with two replicate per donor-timepoint value, from [29]. Figure 2 displays plots of the first two coordinates of each replicate grouped by donor and timepoint. We see that for almost all summaries, these points cluster according to donor identity, with the CDR3 pairwise distance distribution being the only summary that does not decisively cluster by donor. Many summaries additionally cluster according to timepoint in the second dimension, although the tightness of clustering varies by summary, with some summaries (e.g. DJ insertion length distribution) being tightly clustered by a given donor/timepoint value and some summaries (e.g. Kidera factor 4) not obviously clustering by donor/timepoint. Moreover, the D gene usage distribution for each individual splits into two distinct groups which do not correlate with timepoint, though the import of this is more difficult to assess. Although these patterns would require further exploration in a particular research context, these `sumrep` divergences show interesting patterns when TCR datasets are stratified by covariates.

**Figure 2:**
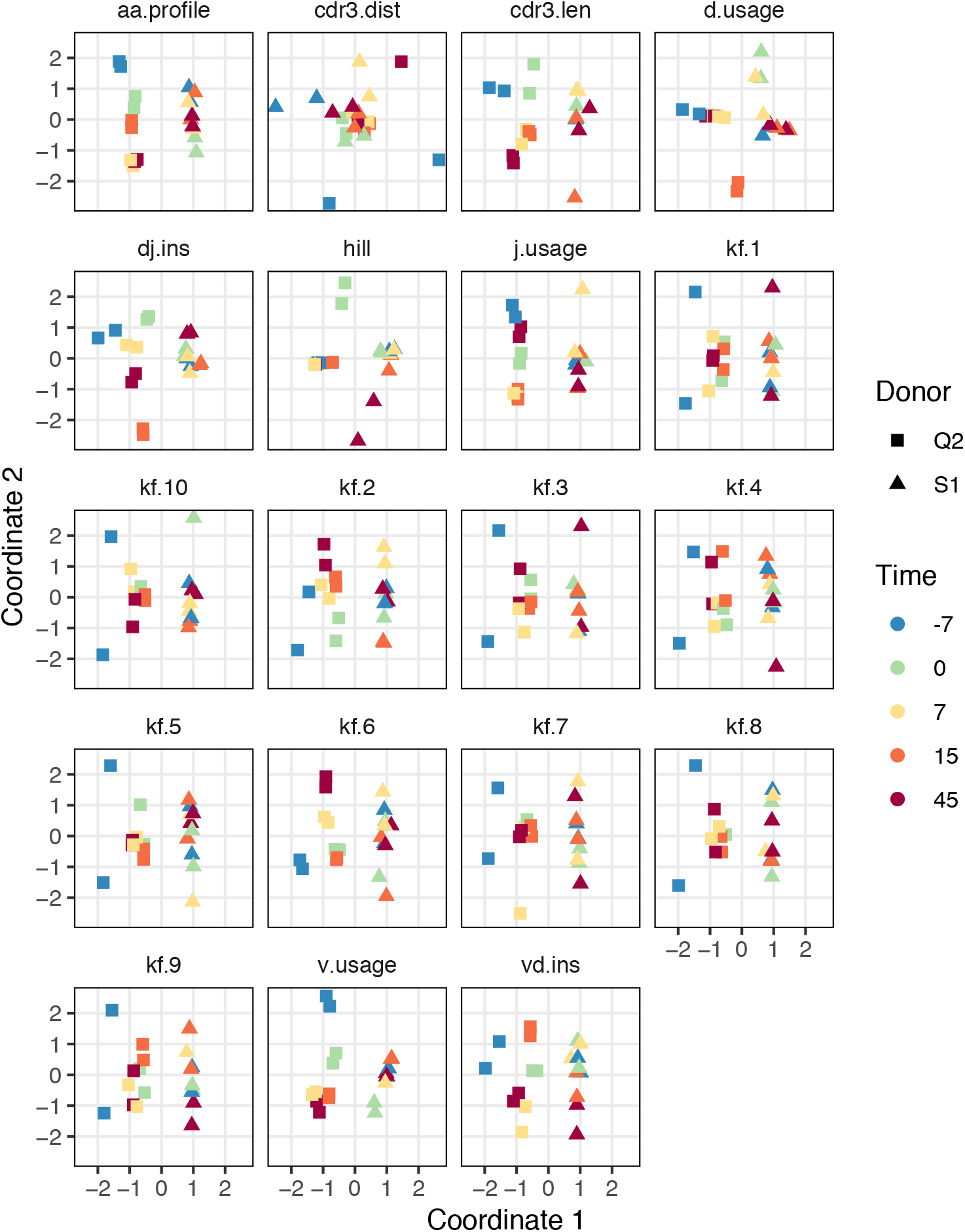
Plots of summary divergence MDS coordinates for data from Pogorelyy et al, 2018, grouped by donor and timepoint

We performed a similar MDS analysis of summary divergences of BCR repertoires stratified by covariate, using data from [32]. We computed divergences of each summary between each pair of a collection of datasets stratified by five pairs of twins as well as B cell classification as memory or naive to form a dissimilarity matrix. We then mapped these dissimilarity matrices to an abstract Cartesian space using MDS. Figure 3 displays plots of the first two coordinates of each donor grouped by twin pair identity and cell type. We see that for each summary, points can be separated according to cell subset, with some summaries (e.g. V gene usage, AA frequencies, acidity) clustering more tightly among cell subset, and others (e.g. GRAVY index, DJ insertion length) clustering more loosely. In addition, the naive repertoires appear to be more tightly clustered than the memory repertoires for each summary. Finally, for the gene usage statistics, there is a strong tendency for twins to have higher similarity than unrelated donors, although this tendency is not consistently observed for other statistics. For example, points for the amino acid 2mer frequency distribution divergences tend to have high similarity between twins, but the GRAVY index distribution divergences do not. Thus, there seem to also be interesting dynamics underlying `sumrep` divergences when BCR datasets are stratified by covariates, and the observed patterns merit further investigation.

**Figure 3:**
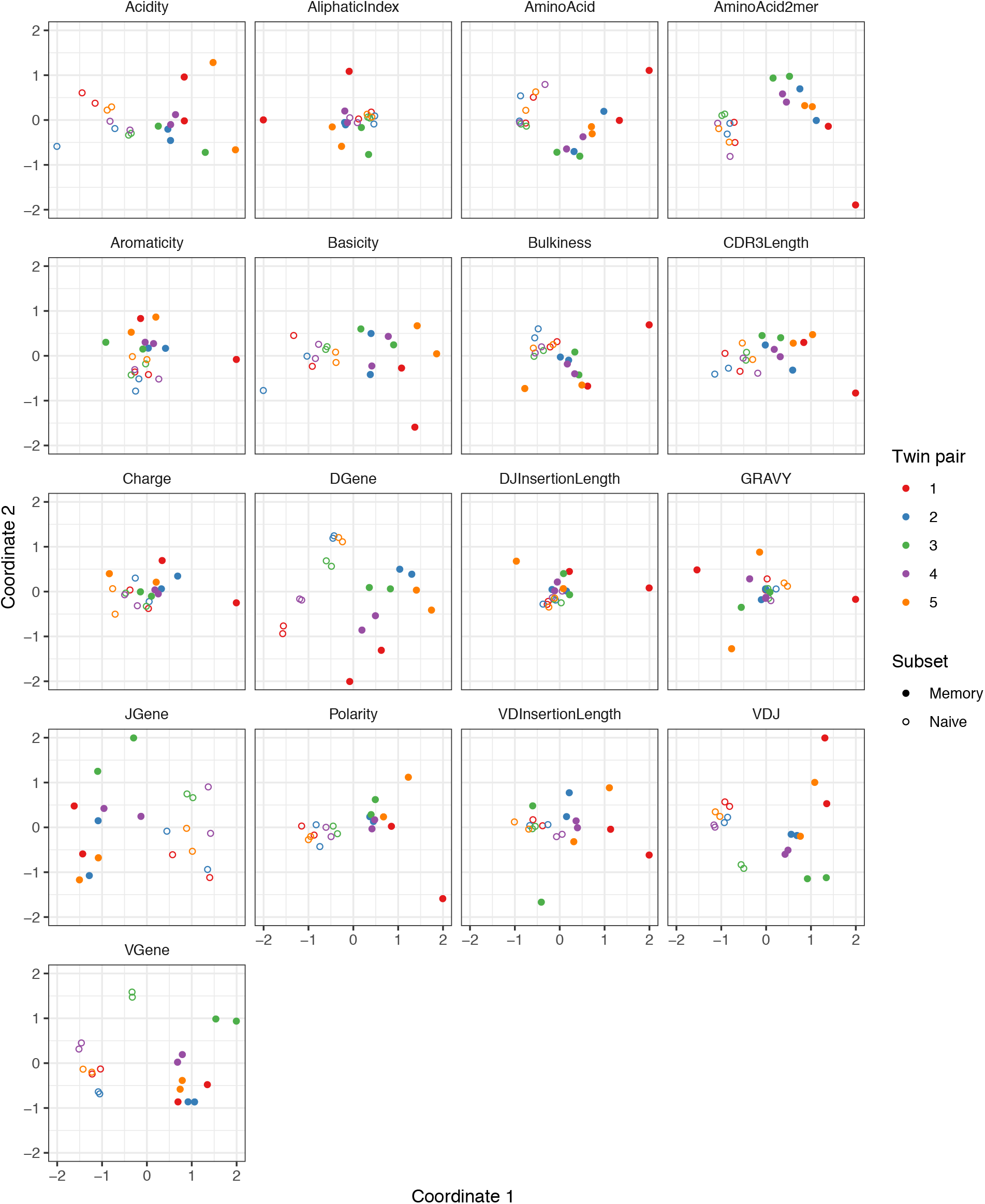
Plots of summary divergence MDS coordinates for data from Rubelt et al, 2016, grouped by twin pair identity and cell type (memory vs naive).

### Ranking summary statistic informativeness

Due to the large number of summary statistics supported by `sumrep`, many of which are correlated, we sought an approach to identify a set of maximally-informative statistics that provide complimentary information to one another. To address this, we employed a lasso multinomial regression treating certain sequence-level summaries as covariates and dataset identity as the response. The basic idea is that this regression method cuts out all but a few predictor variables to find a smaller collection of informative summary statistics, as a coefficient is “allowed” to be nonzero only when the lasso deems it a relatively meaningful predictor. As the regularization parameter *λ* is decreased, more and more coefficients become nonzero, leading to a natural ordering of summaries as the order in which their coefficient “branches off” from zero. Then a resultant maximally-informative set of *k* summaries is the set of summaries with the *k* best ranks. We formalize this approach in the Methods section (Algorithm 3).

One caveat to this approach is that we can only use sequence-level summary statistics as covariates in order to have a well-defined regression procedure. However, the majority of summaries considered in this report are applied at the sequence level. Thus, between the subset of informative sequence-level statistics and the remaining non-sequence-level statistics, we arrive at a considerably smaller set. Besides non-sequence-level summaries, we also omit Kidera Factors and Atchley factors from our analyses as these sets of statistics are orthogonal by construction according to particular measures of amino acid composition in their respective original contexts. This also leads to a much smaller design matrix and a substantially decreased runtime.

Figure 4a displays the results of applying Algorithm 3 to `IGoR` annotations of TRB sequences from datasets A4 i107, A4 i194, A5 S9, A5 S10, A5 S15, and A5 S22 from Britanova et al, 2016 [6]. We see that recombination-based deletion lengths comprise four of the top five summaries, with recombination-based insertion lengths, CDR3 length, and various physiochemical CDR3 properties scattered over the remaining positions. There appears to be high variability throughout the range of rankings, with the bottom three statistics all having a ranking of one for at least one coefficient vector.

**Figure 4:**
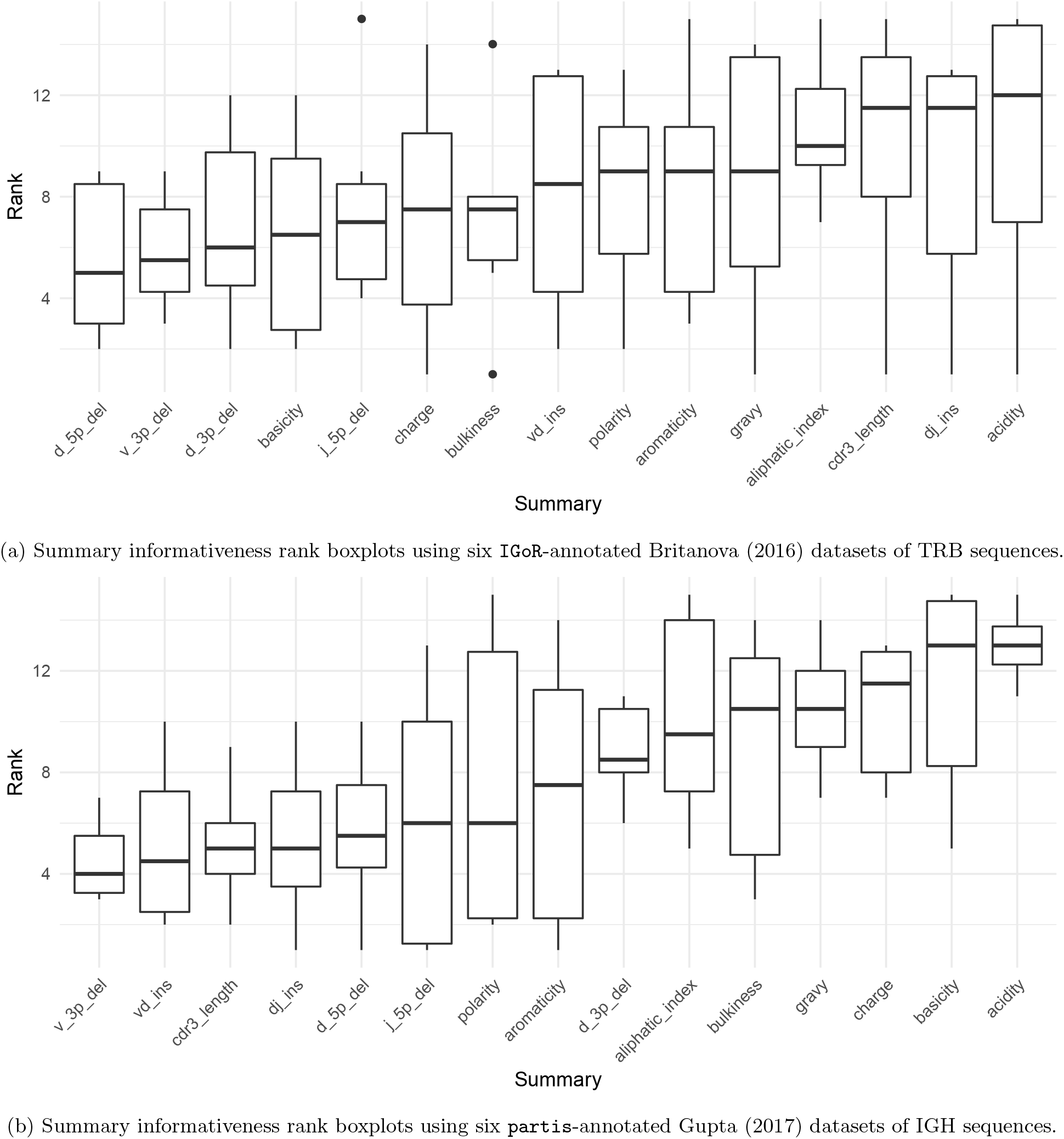
Boxplots of summary rank values taken over each dataset, in order of informativeness, as determined by the median order in which the summary branches off from the lasso paths in Figure S8, taken over each of the six paths.

Figure 4b displays the results of applying Algorithm 3 to `partis` annotations of IGH sequences from donors FV, GMC, and IB at timepoints −8d and −1h from Gupta et al, 2017 [12], downsampled to unique clonal families to avoid clonal abundance biases and decrease algorithmic runtime. We see that deletion lengths, insertion lengths, and CDR3 length comprise the top six summaries, with physiochemical CDR3 properties mostly in the bottom half of rankings. In contrast to the TCR result, there appears to be less overall variability throughout the range of rankings, with variability highest for the moderate ranking positions and notably lower for the top and bottom positions.

While it’s difficult to say exactly the level of correlation of each summary by the lasso result alone, since the lasso is a regularized version of least-squares, our intuition is that the nice properties of least-squares combined with the lasso’s ability to eliminate less relevant coefficients leads to a subset of covariates that are generally informative. To validate this intuition, we can examine distributions of particularly ranked summaries applied to a test set of annotated repertoires not used in the model fitting. Figure 5 displays ECDFs of the the acidity (bottom-ranked), aromaticity (middle-ranked), and V 3’ deletion length (top-ranked) distributions for the FV, GMC, and IB donors at timepoints +1h, +7d, and +28d following an influenza vaccination (which differ from the −1h and −8d timepoints used for fitting), where the ranks are as determined by Figure 4b for `partis`-annotated IGH repertoires. Visually, we see that the acidity curves do not vary much among donors or timepoints; the aromaticity curves have slightly more variation but are still highly similar; and the V 3’ deletion length curves are more distinguished between some donors (e.g. FV and GMC) as well as some donor-timepoint interactions (e.g. +7d and +28d timepoints for IB). Thus, there is visual evidence that the lasso scores can identify some degree of informativeness among summaries.

**Figure 5:**
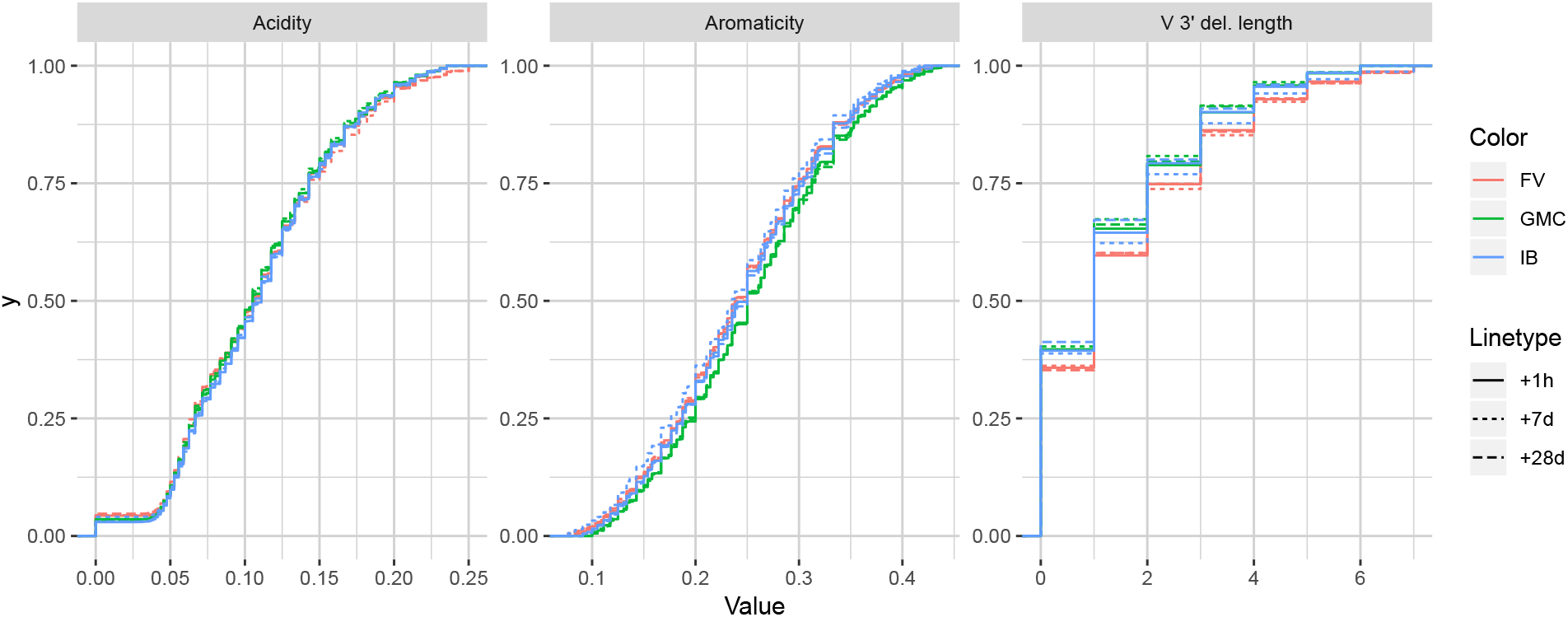
Empirical cumulative distribution functions for the bottom-, middle-, and top-ranked statistics for `partis`-annotated IGH repertoires, as determined by Figure 4b.

### Comparing experimental observations to model simulations

`sumrep` can be used to validate BCR/TCR generative models, i.e. models from which one can generate (simulate) data, through the following approach. First, given a collection of AIRR-seq datasets, model parameters are inferred using the modeling software tool for each repertoire, and then these parameters are used to generate corresponding simulated datasets (Figure 1c). Next, `sumrep` is used to compute the summary statistics listed in Table 1 for each dataset and compare these summaries between each pair of datasets (Figure 1d). Then, a score is calculated for how well the software’s simulation replicates a given summary based on how small the divergences of observed/simulated dataset pairs are compared to divergences between arbitrary observed/observed or simulated/simulated pairs.

Applying this methodology using many datasets should give a clear view of which characteristics the model captures well, as well as areas for improvement. As described in the introduction, we are motivated to do this because models are often benchmarked on simulated data, and it is important to understand discrepancies between simulated and observed data in order to properly interpret and extrapolate benchmarking results. We emphasize that validating the model in this way is different than the usual means of benchmarking model performance: rather than benchmarking the inferential results of the model, we benchmark the model’s ability to generate realistic sequences.

We illustrate this approach with two case studies: an analysis of `IGoR` [20] applied to TRB sequences, and an analysis of `partis` [30, 31] simulations applied to IGH sequences. Both tools are applied to separate sets of experimental repertoires, yielding model-based annotations for each repertoire, as well as simulated datasets from the inferred model parameters for each experimental set. Summary divergences are applied to each dataset, allowing for scores for each summary to be computed for each tool.

### Assessing summary statistic replication for IGoR

We apply the methodology discussed in the previous section to TRB sequences from datasets A4_i107, A4_i194, A5_S9, A5_S10, A5_S15, and A5_S22 from Britanova et al, 2016 [6]. Although `IGoR` is typically applied to non-productive sequences in order to capture the pre-selection recombination process, for this example application we wished to understand `IGoR`’s ability to fit the complete repertoire directly without the need for an additional selection model (e.g. [9]). Thus, we fit the `IGoR` model with all sequences (which we expect to be dominated by productive sequences) and restricted the simulation to productive sequences. Figure 6 contains frequency polygons of each summary distribution for each experimental and simulated repertoire.

**Figure 6:**
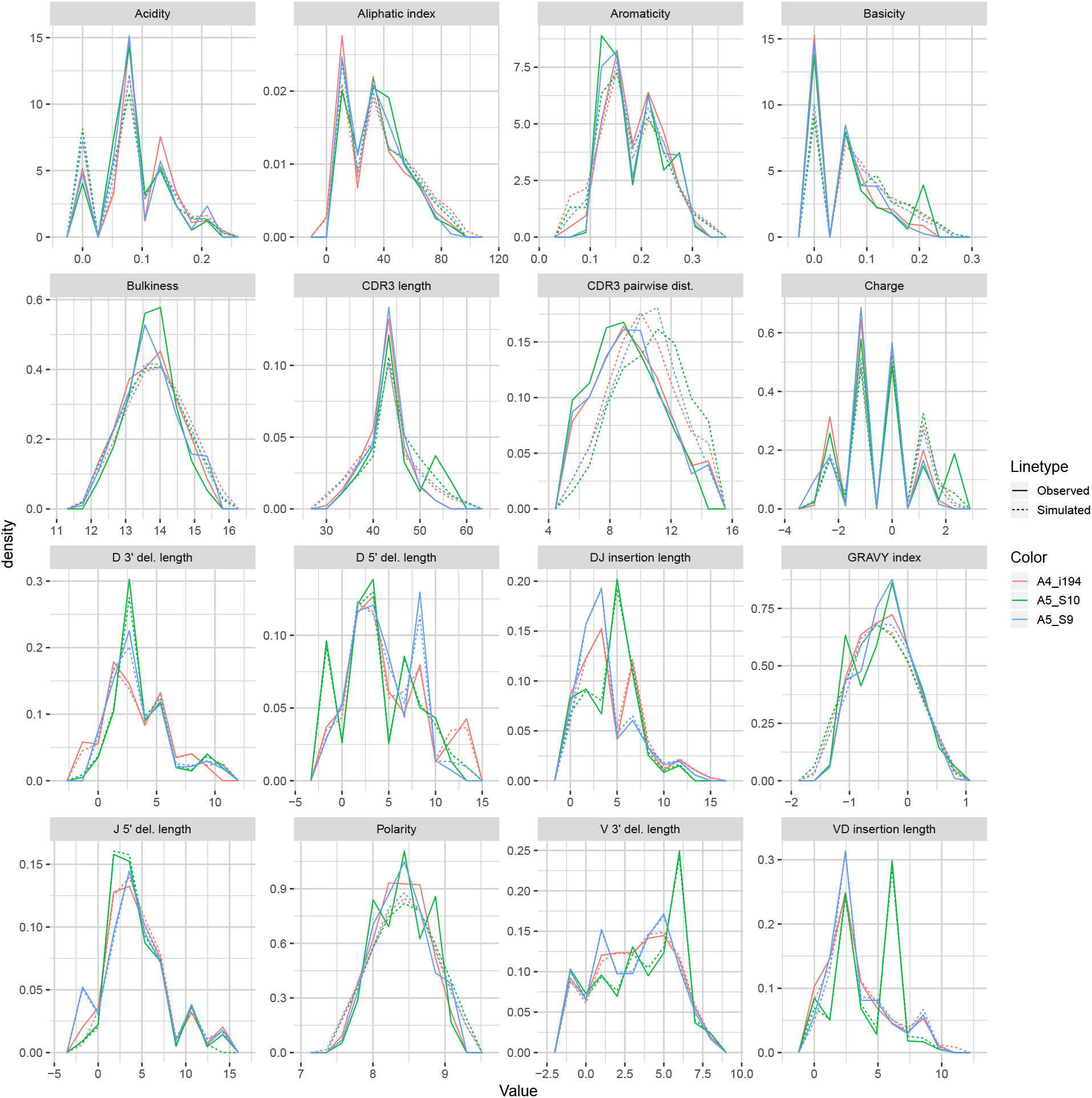
Frequency polygon plots of each univariate summary distribution for the IGoR datasets.

Observation-based summary scores are computed using a log ratio of average divergences (referred to as LRAD-data, and defined in (8)) for a variety of TRB-relevant summaries (Figure 7a). The LRAD-data score of a summary will be high when simulations look like their corresponding observations with respect to that summary, and low when observations look more like other observations than their corresponding simulations. We exclude summaries based on `sequence_alignment` values (e.g. pairwise distance distributions) since `IGoR` does not currently have an option to output the full variable region nucleotide sequences for experimental reads.

**Figure 7:**
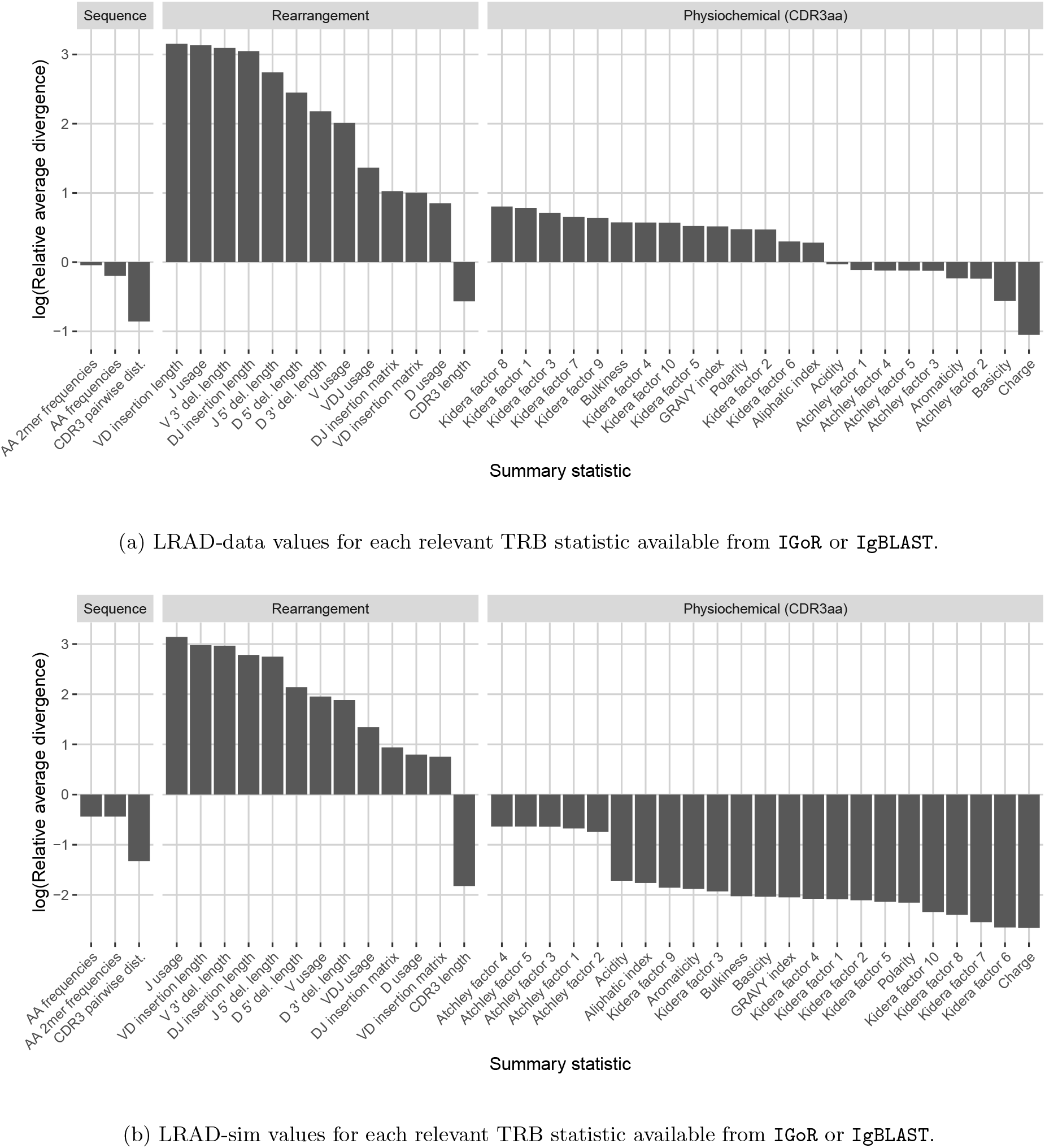
Summary scores, denoted as “log(Relative average divergence)” or “LRAD,” for each statistic in the `IGoR` model validation experiment. For both cases, a high score indicates a well-replicated statistic by the simulations with respect to their corresponding experimental repertoires of functional TRB sequences.

`IGoR` simulations were able to recapitulate gene usage statistics of an empirical repertoire well, with J gene usage frequency being the most accurately replicated, followed by various recombination-based indel statistics. V, D, and joint VDJ gene usage are all also well-replicated, as well as both VD and DJ insertion matrices. Conversely, the CDR3 length distribution was the least accurately replicated statistic among rearrangement statistics. The Kidera factors of the CDR3 region were also replicated well, despite CDR3 length being one of the least accurately replicated statistics. Scores for other CDR3-based statistics besides Kidera factors ranged from mildly good to mildly bad, with the GRAVY index distribution being the best CDR3-based statistic (excluding Kidera factors) and charge distribution being the worst.

We also computed simulation-based summary scores (LRAD-sim, defined in (9)) for the same datasets and simulations (Figure 7b). The LRAD-sim score of a summary will be high when simulations look like their corresponding observations with respect to that summary, and low when simulations look more like other simulations than their corresponding observations. We still saw high scores for gene usage and indel statistics, although the CDR3 length distribution and various Kidera factor and GRAVY index distributions had much lower scores. This suggests that while the average `IGoR` simulation yields Kidera factor and GRAVY index distributions that look more like the observed repertoire’s distributions than other observed repertoires do, these simulated repertoires still tend to produce more similar distributions to each other than to their observed counterparts. In turn, this provides an avenue of future research for TCR generative models in which certain CDR3aa properties are incorporated and expressed in simulated data.

### Assessing summary statistic replication for partis

We applied the same methodology to IGH sequences from Gupta et al, 2017 [12], using datasets corresponding to the −1h and −8d timepoints for each of the FV, GMC, and IB donors. Figure 8 displays frequency polygons of each summary distribution for each experimental and simulated repertoire.

**Figure 8:**
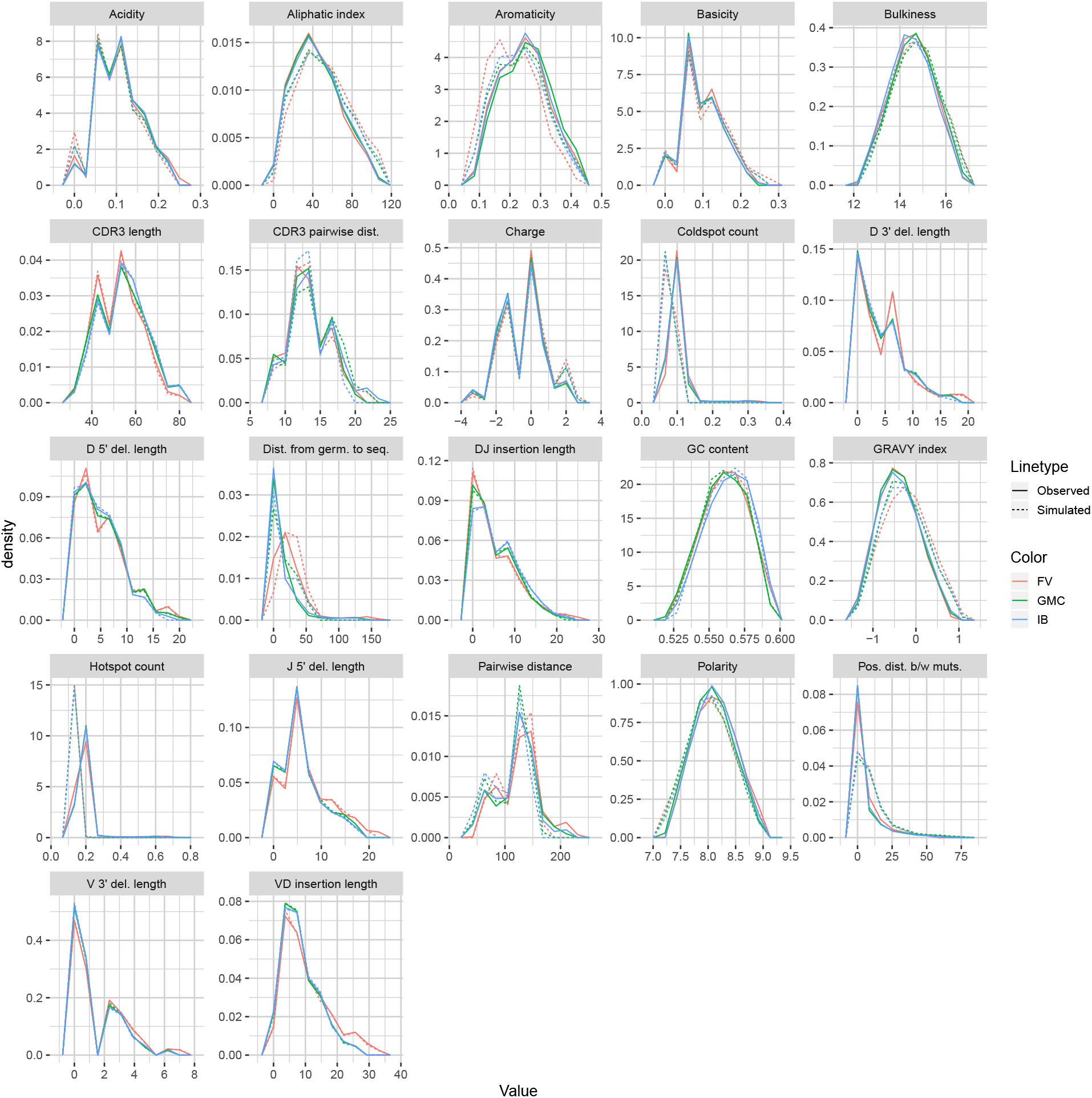
Frequency polygon plots of each univariate summary distribution for the `p_f1`, `p_f1_sim`, `p_g1`, and `p_g1_sim` datasets.

Observation-based summary scores were computed using the LRAD-data equation (8) for a variety of IGH relevant summaries (Figure 9a).

**Figure 9:**
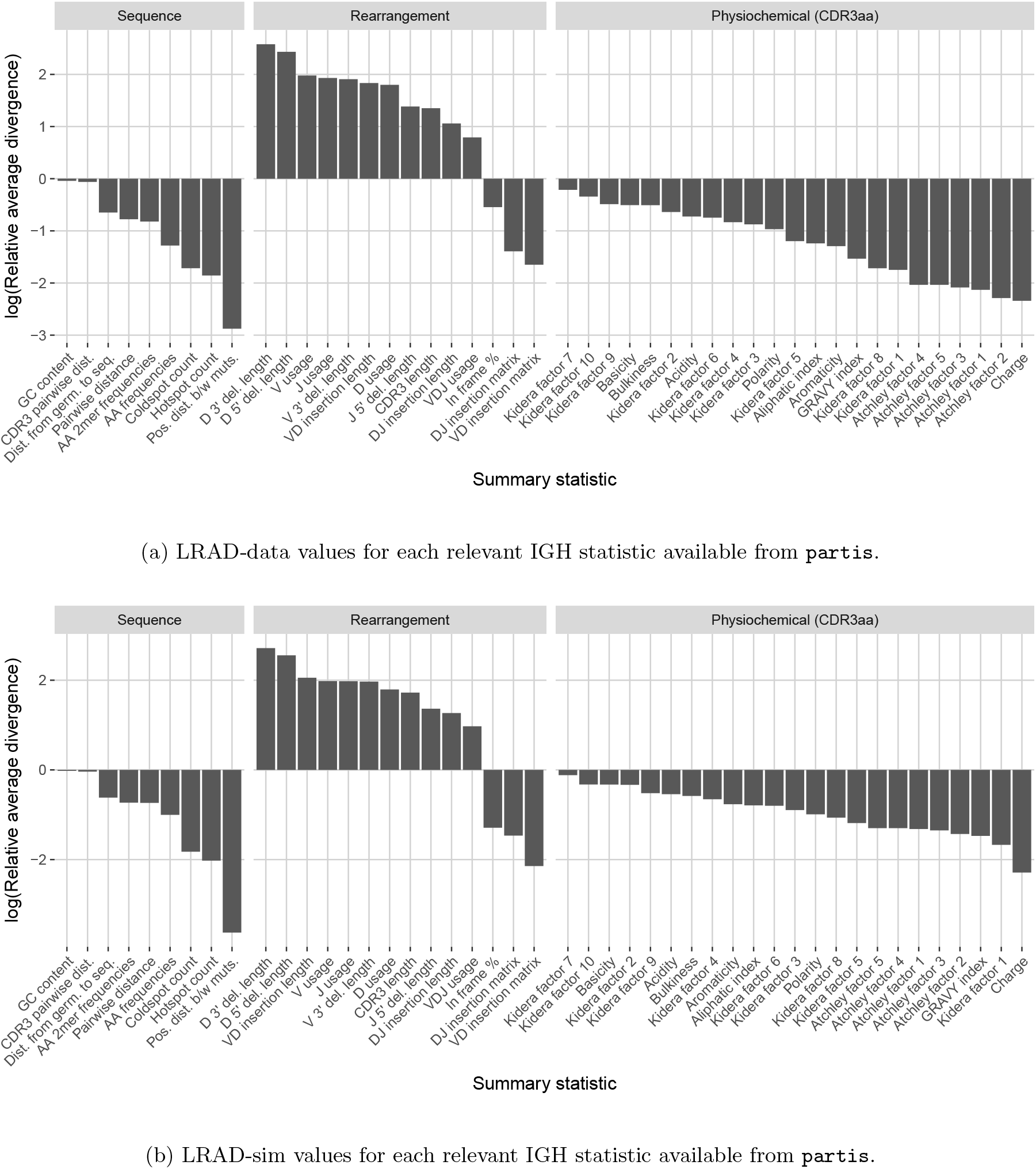
Summary scores, denoted as “log(Relative average divergence)” or “LRAD,” for each statistic in the `partis` model validation experiment. For both cases, a high score indicates a well-replicated statistic by the simulations with respect to their corresponding experimental repertoires of productive IGH sequences.

Like `IGoR`, we see that `partis` simulations also excelled at replicating gene usage and recombination statistics, while additionally replicating CDR3 length distributions well. However, `partis` struggled to recapitulate VD and DJ insertion matrices, which it does not explicitly include in its model. This contrasts with `IGoR` which incorporates these insertion matrices during model fitting, and thus recapitulates these matrices well. The other statistics yielded scores ranging from slightly to very negative, with many mutation-based summaries like positional distance between mutations and hot and cold spot counts being poorly captured. The low scores of mutation-based summaries may arise from the decision to select a single representative from each clonal family, which itself arises from the complications in matching clonal family abundance distributions of simulations to data. This makes it difficult to identify the exact contributions of these factors to the summary discrepancies. Nonetheless, this suggests that these sorts of quantities may need to be more explicitly accounted for in BCR generative models if more realistic simulations are desired.

We also computed simulation-based summary scores (LRAD-sim, defined in (9)) for the same datasets and simulations (Figure 9b). The scores are highly similar to those seen in Figure 9a, with some summaries seeing a moderate drop.

## Methods

### Divergence

We use the Jenson-Shannon (JS) divergence for comparing distributions of scalar quantities, which constitutes most summaries in `sumrep`. The Jenson-Shannon divergence of probability distributions *P* and *Q* with densities *p*(*⋅*) and *q*(*⋅*) is a symmetrized Kullbeck-Leiber divergence, defined as

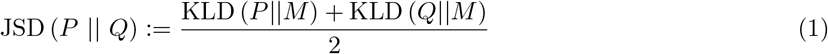

where *M*:= (*P* + *Q*)/2 and KLD(*P ||M*) is the usual KL-divergence,

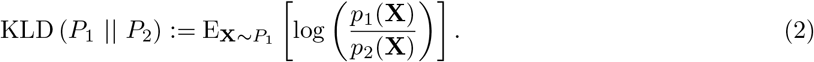

In the case where *P* and *Q* are both discrete distributions, this becomes

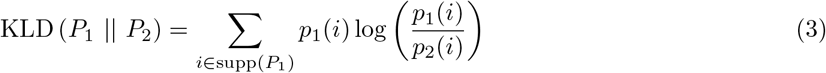

where supp(*P*) is the countable support of distribution *P*. Because the discrete formulation has computational benefits over the continuous one, we discretize continuous samples and treat them as discrete data. By default, we use 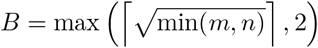 bins of equal length, where *m* = |supp(*P*)| and *n* = |supp(*Q*)|, which is designed to scale with the complexity of *m* and *n* simultaneously. We also discard bins which would lead to an infinite KL divergence for numerical stability.

For counts of categorical data, we instead appeal to the sum of absolute differences, or ℓ_1_ divergence, for comparison:

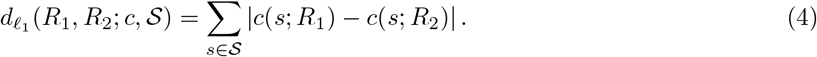

In words, (4) iterates over each element *s* in some set 𝒮, calculates the count *c* of *s* within repertoires *R*_1_ and *R*_2_ respectively, takes the absolute difference of counts, and appends this to a rolling sum. This metric is well suited for comparing marginal or joint V/D/J-gene usage distributions. For example, if 𝒱, 𝒟, and 𝒥 represent the germline sets of V, D, and J genes, respectively, define usage *u* of gene triple (*v, d, j*) ∈ 𝒱 × 𝒟 × 𝒥 for repertoire *R* as

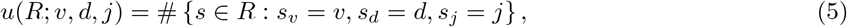

where e.g. *s*_*v*_ = the V gene of *s*. Then an appropriate divergence for the joint VDJ gene usage for repertoires *R*_1_ and *R*_2_ is

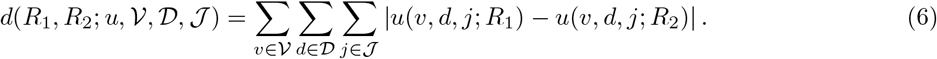

The ℓ_1_ divergence is also relevant for computing amino acid frequency and 2mer frequency distributions. Note that we can normalize the counts to become relative frequencies and apply (4) on the resultant scale which may be better suited to the application, especially when dataset sizes differ notably.

### Approximating distributions via subsampling and averaging

Computing full summary distributions over large datasets can be intractable. However, we can compute a Monte Carlo distribution estimate by repeatedly subsampling and aggregating summary values until convergence. Algorithm 1 formalizes this idea, appending batch samples of *d* to a rolling approximate distribution and terminating when successive distribution iterates have a JS divergence smaller than tolerance *ε*. Note that continually appending values to a rolling vector is analogous to computing a rolling average, where the subject of the averaging is an empirical distribution rather than a scalar.

#### Algorithm 1

Compute automatic approximate distribution

**Input:** repertoire *R*, summary *s*, batch size *m*, convergence tolerance *ε*

**Output:** subsampled approximation to *d*

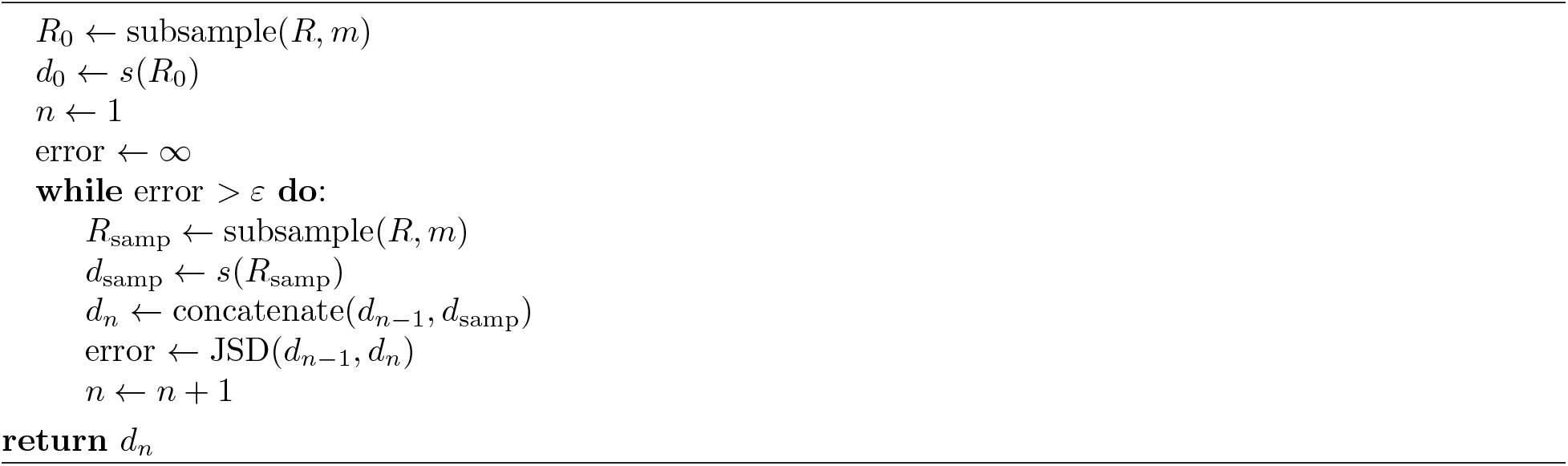

An alternative would be to simply compute the distribution on one subsample of the data and use this as a proxy distribution. The main advantage of Algorithm 1 over such an approach is that it provides a sense of convergence to the full distribution via the tuning parameter *ε*, while automatically determining the size of the necessary subsample. The algorithm can also be tuned according to batch size *m*, which `sumrep` takes to be 30 by default. We conduct a performance analysis of Algorithm 1 in Appendix A and empirically demonstrate efficiency gains in a variety of realistic settings without sacrificing much accuracy.

Some summaries induce distributions for which Algorithm 1 is inherently ill-suited. This occurs when a summary applied to a subset of a dataset does not follow the same distribution as the summary applied to the full dataset. For example, consider the nearest neighbor distance of a sequence *s*_*i*_ with respect to a multiset of sequences *R* (i.e. elements in *R* can have multiplicity *≥* 1),

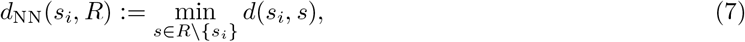

where *d*(*⋅, ⋅*) is a string metric (e.g. the Levenshtein distance). If we take any subset *S* of *R*, then *d*_NN_(*s*_*i*_, *S*) *≥ d*_NN_(*s*_*i*_, *R*) ∀*i*, since *R* will have the same sequences to iterate over, and possibly more sequences, which can only result in the same or a smaller minimum.

In this case, we can still obtain an unbiased approximate to the nearest neighbor distance distribution using the following modification of Algorithm 1. For each iteration, sample a small batch *B* = (*s*_1_,…, *s*_*b*_) of *b* sequences, and compute *d*_NN_ of each *s*_*i*_ to the full repertoire *R*. Since each batch *B* computes the exact nearest neighbor with respect to *R*, we get the true value of *d*_NN_ for each *s ∈ B*. The gain in efficiency stems from the fact that we only compute this true *d*_NN_ for a subsample of the sequences of the full repertoire *R*. Thus, appending batches to a running distribution until convergence as in Algorithm 1 will produce increasingly refined, unbiased approximations as the tolerance decreases. Algorithm 2 explicates this procedure.

Algorithm 2 may yield a high runtime if *R* is large, the sequences in *R* are long, or the tolerance is small. Nonetheless, we empirically demonstrate in Appendix B that in the case of typical BCR sequence reads, even very small tolerances incur reasonable runtimes, and when *R* is large, the algorithm is orders of magnitude faster than computing the full distribution over *R*.

#### Algorithm 2

Compute automatic approximate nearest neighbor distance distribution

**Input:** repertoire *R*, distance *d*, batch size *m*, convergence tolerance *ε*

**Output:** subsampled approximation to *d*_NN_

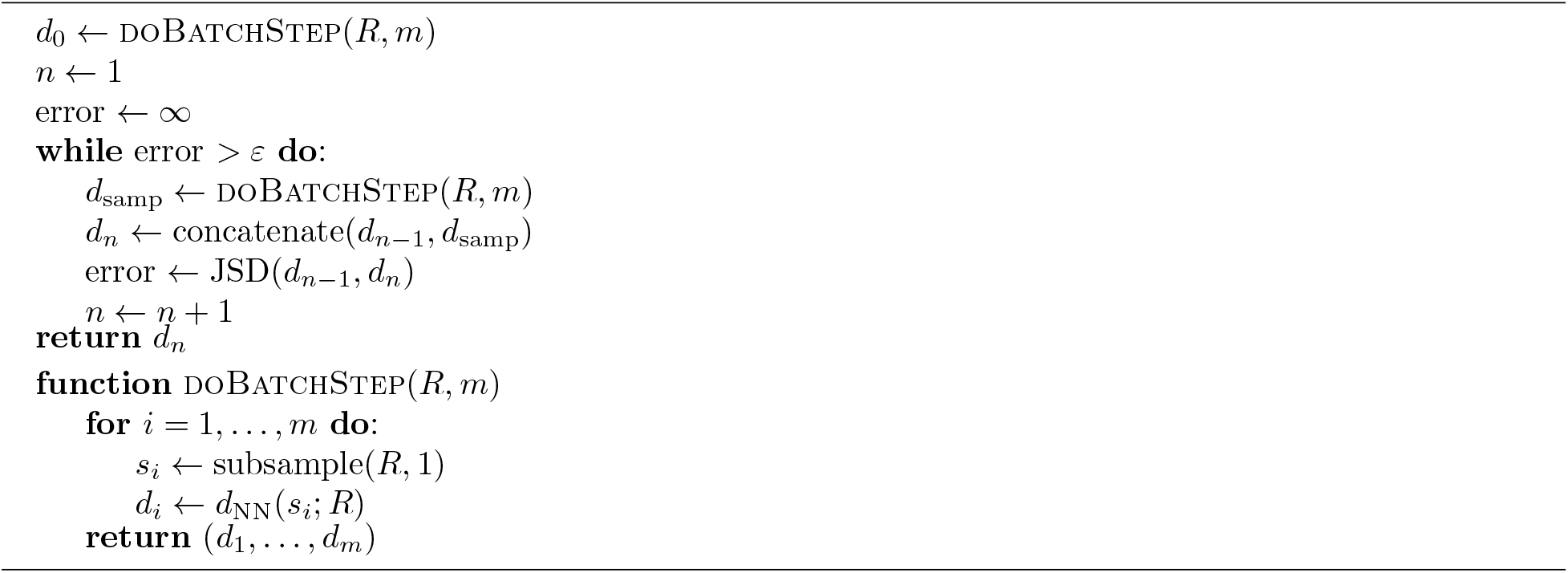

### Summary statistic informativeness ranking

To quantify the relative informativeness of various summary statistics in distinguishing between different datasets, we perform a multinomial lasso regression where covariates are sequence-level summaries and the response is dataset identity. Since ℓ_1_ multinomial regression outputs a separate coefficient vector ***β*** for each response value, we aggregate by taking medians of each dataset-specific lasso ordering for each summary to get the final score. This also yields a range of rankings to assess the variation in scores by summary and by inferential model (e.g. `partis`, `IGoR`). In the case of ties, we randomize rankings to avoid alphabetization biases or other similar artifacts. Detailed pseudocode is provided in Algorithm 3.

#### Algorithm 3

Rank summary statistics by informativeness

**Input:** annotations datasets *d*_1_,…, *d*_*D*_, sequence-level summaries **s**(*⋅*) = [*s*_1_(*⋅*),…, *s*_*S*_ (*⋅*)], lasso parameters *λ*_1_,…, *λ*_Λ_

**Output:** A vector of ranks for the summaries

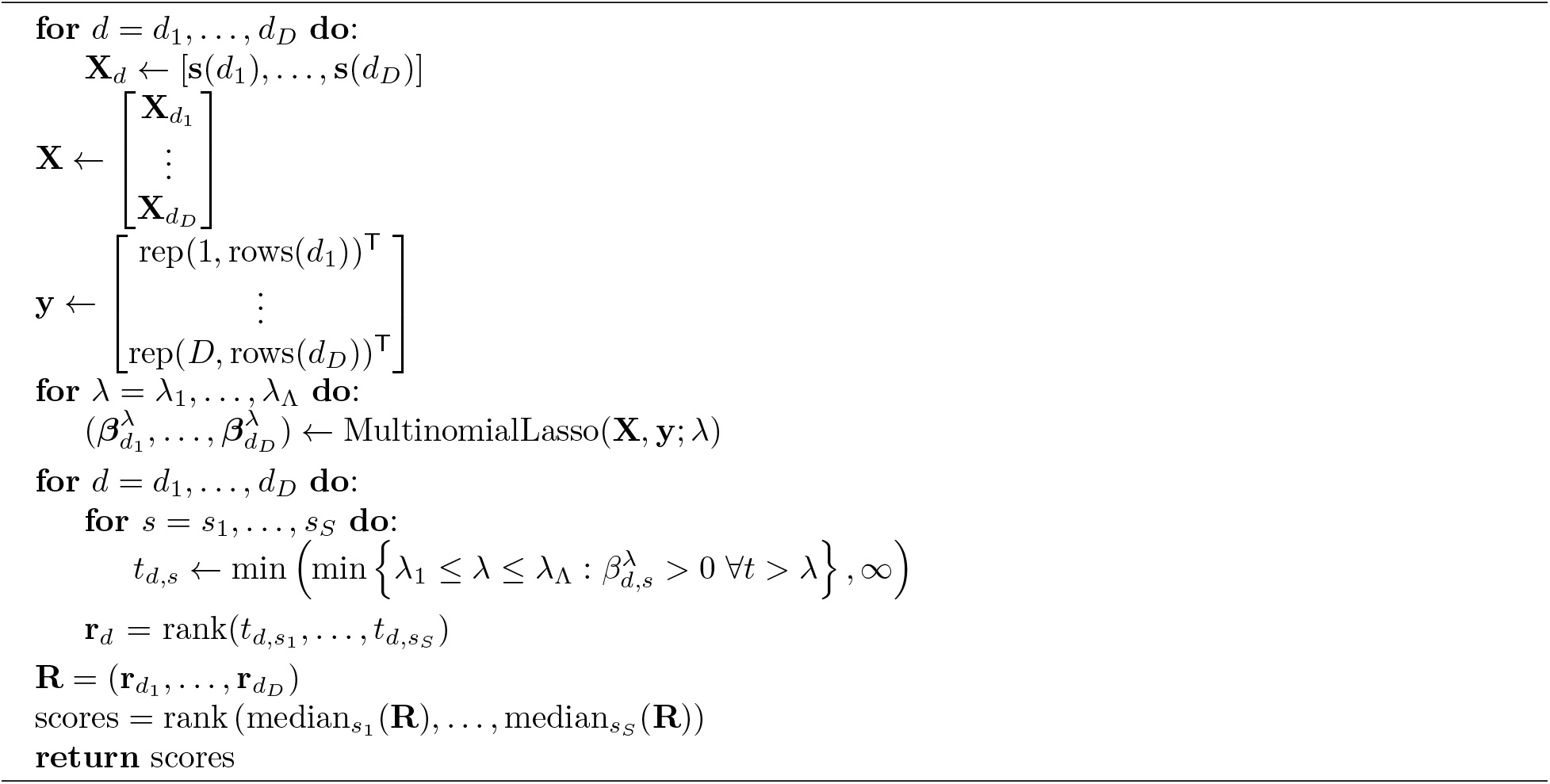

This approach only works for sequence-level summaries *s ∈* ℝ^*n*^ for a dataset *d* of *n* = rows(*d*) sequences in order to form a well-defined design matrix 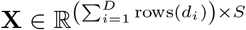 over all datasets *d* = *d*_1_,…, *d*_*D*_ under consideration. For example, it is unclear how to incorporate the pairwise distance distribution, which is not a sequence-level summary, as a covariate, since this summary in general yields a column of a larger length than the number of sequences. Still, as most summaries considered above can be applied at the sequence level, this method greatly reduces the number of summaries the user needs to examine.

### Model validation of IGoR

We used the `-infer` subcommand of `IGoR` to fit custom, dataset-specific models for each experimental dataset. Since we were interested in many CDR3-based statistics and `IGoR` does not currently include inferred CDR3 sequences with rearrangement scenarios, we used `IgBLAST` to extract CDR3s for each sequence. For each sequence, we considered only the rearrangement scenario with the highest likelihood as determined by `IGoR`. When a list of more than one potential genes was given as the gene call, we considered only the first gene in the list. Several fields were renamed to match the AIRR specification when the definitions align without ambiguity. As described in Results, we trained on productive sequences and restricted the simulation to productive sequences.

We applied `IGoR` in this way to six datasets of TRB sequences from [6], which studied T cell repertoires from donors ranging from newborn children to centenarians.

### Model validation of partis

We used `partis` to infer custom generative models for each experimental dataset. We ran the `partition` subcommand to incorporate underlying clonal family clustering among sequences during inference, and then downsampled each observed and simulated dataset so that each clonal family is represented by one sequence. Since `partis` returns a list of the top most likely annotations scenarios for each rearrangement event, we considered only the scenario with the highest model likelihood for each sequence. We denote the `indel_reversed_seqs` field as `sequence_alignment` and naive seq as `germline_alignment` as they satisfy these definitions from the AIRR Rearrangement schema. Several other fields are renamed to match the AIRR specification when the definitions align without ambiguity.

Before running summary comparisons, we randomly downsample to one receptor per clonal family to get a dataset consisting of unique clonotypes for both the observed and simulated datasets. We do this since `partis simulate` draws from distributions over clonal families for each rearrangement event as inferred from `partis partition`. While it is possible to simulate multiple leaves for each rearrangement, it is not obvious how to best synchronize this with the observed clonal family distributions. A more involved analysis would attempt to mimic the clone size distribution in data as closely as possible, potentially with correlations between clone size and other rearrangement parameters, and assess sequence-level statistics within each clonal family. Here we opt to subsample to unique clones and avoid abundance biases altogether.

We applied `partis` in this way to six datasets of IgH sequences from [12], which studied B cell repertoires from donors prior to and following an influenza vaccination.

### Scoring summary statistic replication by model

We wish to measure how well a given statistic is replicated when a model performs simulations using parameters inferred from an observed repertoire dataset. One approach is to score the statistic *s* based on the average divergence of observations to their simulated counterparts when applying *s*(*⋅*), and the average divergence of observations to other observations when applying *s*(*⋅*). Suppose we have *k* experimental repertoires of immune receptor sequences, and let *R*_*i,obs*_ and *R*_*i,sim*_, 1 *≤ i ≤ k*, denote the *i*th observed and simulated repertoire, respectively. For a given statistic *s*, let 𝒟_*s*_(*R*_1_, *R*_2_) be the divergence of repertoires *R*_1_ and *R*_2_ with respect to *s*. We can score a simulator’s ability to recapitulate *s* from the observed repertoire to the simulated via the following log relative average divergence (LRAD):

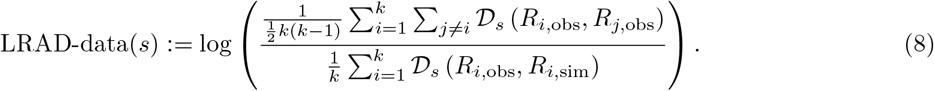

For a given summary *s*, LRAD-data will be positive if the simulated repertoires tend to look more like their experimental counterparts in terms of this summary than experimental repertoires look like other experimental repertoires, and negative if experimental repertoires tend to look more like other experimental repertoires than they do their simulated counterparts. In other words, LRAD-data scores how well a simulator can differentiate *s* from an experimental repertoire among other repertoires, and recapitulate *s* into its simulation. Applying the log to the ratio allows for the magnitudes of scores to be directly comparable (so that a summary with score −*a >* 0 performs as well as a summary with score *a <* 0 performs poorly).

Another related score compares the average divergence of observations to their simulated counterparts, and the average divergence of simulations to other simulations. Formally, this becomes

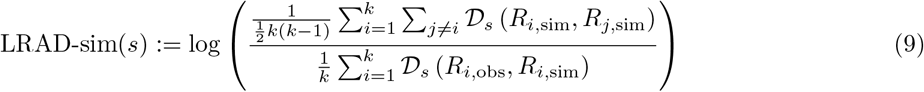

where the difference from (8) is that the divergences in the numerator are applied to simulated-simulated dataset pairs rather than observed-observed dataset pairs. LRAD-sim for a given summary will be positive if simulated repertoires tend to look more like their experimental counterparts in terms of this summary than simulated repertoires look like other simulated repertoires, and negative if the simulated repertoires tend to look more alike.

These scores underlie the model validation analyses of `partis` and `IGoR` simulations in the Results section, and comprise the values displayed in Figures 7 and 9. However, this framework can be used to validate any immune receptor repertoire simulator which outputs the fields compatible with the summaries in Table 1, or more generally any set of summaries generated by a model-based simulator that is not supported directly by `sumrep`.

A feature of our methodology is that we use the same tool to produce simulations that we used to produce the annotations. To examine the sensitivity of this method, we performed a separate analysis by obtaining dataset annotations from standalone `IgBLAST` [38], and comparing these to simulations based on `partis` annotations using IMGT germline databases. We did not perform a similar analysis for `IGoR` annotations since `IgBLAST` was used to infer CDR3s within the `IGoR` workflow. This is discussed in detail in Appendix C.

### Materials

The raw data for the TCR summary divergence MDS analysis comes from [29], which was postprocessed into a suitable format for analysis. For each donor-timepoint combination, a single blood draw was split in replicas at the level of cell mixture.

The raw data for the BCR summary divergence MDS analysis comes from [32]; `IgBLAST`-preprocessed data was downloaded from VDJServer in the AIRR format. For quality control, sequences with a run of 3 or more N bases in the raw sequence were discarded.

For the TCR model validation analysis, we use six datasets from [6], corresponding to labels A4_i107, A4_i194, A5_S9, A5_S10, A5_S15, and A5_S22. For tractability purposes, we chose the six datasets with the fewest number of sequence reads; the number of reads from these six datasets used in the analysis ranged from 37,363 sequences to 243,903 sequences. These datasets consist of consensus RNA sequences assembled using UMIs. Most of these sequences are productive; as previously described, for this example application we are benchmarking `IGoR`’s ability to fit complete repertoires rather than only non-productive repertoires.

The data for the BCR model validation analyses originated from samples first sequenced and published in [18], although we used the Illumina MiSeq data published in [12] for our analyses. These datasets represent repertoires of three human donors from multiple time points following an influenza vaccination. We use datasets from time points −1h and −8d for the FV, GMC, and IB donors for the summary informativeness and `partis` model validation analyses; the +1h, +7d, and +28d datasets for the FV, GMC, and IB donors for the summary informativeness validation; and the FV −1h dataset for the approximation routine performance analyses in appendices 1 and 2.

## Conclusions

We have presented a general framework for efficiently summarizing, comparing, and visualizing AIRR-seq datasets, and applied it to several questions of scientific interest. One can imagine many further applications of `sumrep`, as well as promising avenues of research: contrasting repertoires in the context of antigen response or vaccination design and evaluation may shed some light on which summaries can distinguish between such covariates; and comparing the summary distributions of naive repertoires from multiple healthy individuals is likely to aid our understanding of the patterns of variability exhibited by “normal” repertoires, which in turn may aid the detection of repertoire abnormalities. `sumrep` could also be used to evaluate the extent to which artificial lymphocyte repertoires look like natural ones [10].

There are several other packages dedicated to detailed summaries and visualization of immune receptor repertoires. The `tcR` [25] and `bcRep` [2] packages for R include methods for retrieving and comparing gene usage summaries, computing clonotype diversity indices, and visualizing various repertoire summaries. `VDJtools` [34] is a command line tool which performs similar repertoire summarization, comparison, and visualization tasks for TCR data. Desktop GUI-based programs include `ImmunExplorer` [33] and `Vidjil` [8]. `Vidjil` is also available as a webserver, as is `ASAP` [1]. `Antigen Receptor Galaxy` [16] offers online access to many analysis tools. These tools have a subset of the summary statistics described here, and do not have the comparative analysis features of `sumrep`. The `IGoR` [20] software features an algorithm for summarizing statistics of the V(D)J rearrangement process; however, its main focus is on learning the basic model for non-productive T- and B-cell repertoire and it does not provide any built-in methods for comparing inferred models between datasets.

A natural extension of the model validation in this report would be to assess the performance of many competing repertoire analysis tools over a larger group of datasets. `sumrep` can be also used to detect systemic biases between different library preparation protocols and control for batch effects that can confound meta-analysis of AIRR-Seq data. Moreover, while many of the summaries are applied to the CDR3 region by default, it would be interesting to perform separate analyses restricted to different CDRs and framework regions, as physiochemical characteristics of these regions can differ greatly.

Finally, although `sumrep` already supports the AIRR rearrangement schema by default, we plan to thoroughly integrate `sumrep` as a downstream analysis tool for any AIRR-compliant software or workflow.

## Acknowledgements

We thank Misha Pogorelyy for kindly providing post-processed data from [29] and Quentin Marcou for help running `IGoR`. We also thank the other members of the Software WG for early stage discussion, especially Christian Busse, Enkelejda Miho, Inimary Toby, and Jian Ye.

## Author Contributions

BJO, PM, CS, DKR, JAVH, MS, AS, WL, and FAM conceived of the project and guided the overall design of the software and analyses; BJO designed and implemented the main software package; BJO, PM, CS, and AO performed computational analyses; BJO and FAM wrote the first draft of the manuscript; All authors contributed to manuscript revision, read and approved the submitted version.

## Appendix A Performance analysis of Algorithm 1

Here, we run Algorithm 1 on the `partis`-annotated FV −1h dataset (henceforth referred to as `p_f1`), sub-sampled without replacement to 10,000 sequences for tractability. We compute the pairwise distance distribution of CDR3 sequences for the full subsampled dataset, and approximate distributions with tolerances ε ∈ {0.1, 0.001,…, 10^*−*7^}. We replicate this experiment for 10 trials so that the subsampled dataset remains the same, but a new instance of the subsampling algorithm is run each time. Figure S1a shows a frequency polygon of each distribution and figure S1b shows their empirical cumulative distribution functions. We see that the approximate distributions appear to converge to the full distribution as the tolerance gets smaller. Figure S1c displays the KL-divergence to the true distribution for each tolerance, again indicating convergence to the truth. Figure S1d displays the runtimes and log-runtimes for each tolerance as well as the true “population” runtime for the full dataset; while the runtime grows exponentially as *ε* → 0, the approximation algorithm is still much faster than computing the full distribution for each considered value of *ε*.

Next we investigate the effect of dataset size on the performance of Algorithm 1. For sample sizes *n* ∈ {exp(6),…, exp(10)}, we subsample `p_f1` without replacement to *n* sequences and compute the pairwise distance distribution of CDR3 sequences for the full subsampled dataset as well as those given by tolerances *ε* ∈ {0.1, 0.01,…, 10^*−*5^}. We perform this experiment 10 times for each *n*. Boxplots of the KL-divergence by log(*n*) and tolerance over all trials are displayed in Figure S2a. We see no obvious trend in the effect of dataset size on the KL-divergence for any choice of tolerance for the pairwise distribution. Boxplots of the runtime (in log-seconds) by log(size) and tolerance are shown in Figure S2b, showing that runtime increases with sample size for high tolerance, but tends towards a constant runtime by sample size as tolerance decreases. Boxplots of the log-efficiency by log(size) and tolerance are shown in Figure S2c, where

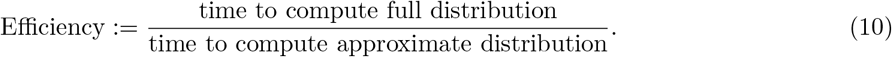

Here we plot efficiency on a log scale, so that the line *y* = 0 corresponds to instances when the true and approximate routines have identical runtimes. Thus, the region *y >* 0 corresponds to instances when Algorithm 1 outperforms the computation of the full nearest neighbor distribution. For moderate to large datasets and reasonable choices of *ε*, the approximate routine is much more efficient than computing the full distribution. Efficiency also appears to increase exponentially with dataset size, although decreases at least exponentially as tolerance decreases. Nonetheless, the accuracy of Algorithm 1 applied to the pairwise distance distribution is scalable to large datasets while leading to large gains in runtime efficiency for reasonable choices of *ε*.

Finally, we investigate the effect of summary statistic on the performance of Algorithm 1. We run the algorithm for the pairwise distance, GC content, hotspot count, coldspot count, and distance from germline to sequence distributions on `p_f1` subsampled without replacement to 10,000 rows. For each summary, we run the algorithm for tolerances *ε* ∈ {0.1,…, 10^*−*5^}. We perform this experiment 10 times for each (summary, *ε*) combination. Figures S3a, S3b, and S3c show the KL-divergence to the full dataset distributions, runtimes, and efficiencies, respectively, by summary and tolerance over all trials. We see that the KL divergence, runtime, and efficiency of the approximation routine depends on the summary in question. In particular, the approximation routine for hotspot and coldspot count distributions does not yield as high of an efficiency for moderately low tolerance, and struggles to minimize the KL-divergence to the true distribution for higher tolerances. This is likely due to the fact that the full hotspot and coldspot count distributions is extremely fast to compute even for large datasets.

These results suggest that convergence and efficiency will vary by summary, and the user should be aware of this fact when choosing whether to run the approximation routine as well as an appropriate tolerance. By default, `sumrep` uses *ε* = 0.001 for arbitrary summary approximation routines, and retrieves approximate distributions by default only for `getPairwiseDistanceDistribution, getNearestNeighborDistribution`, and `getCDR3PairwiseDistanceDistribution`.

## Appendix B Performance analysis of Algorithm 2

Here, we assess the modification of the distribution approximation routine for the nearest neighbor distribution. We run Algorithm 2 on `p_f1` subsampled without replacement to 10,000 sequences for tractability. We compute the nearest neighbor distribution of CDR3 nt sequences for the full subsampled dataset, and approximate distributions with tolerances *ε* ∈ {0.1, 0.001,…, 10^*−*7^}. We replicate this experiment for 10 trials in the same manner as detailed in Appendix A.

Figure S4a shows a frequency polygon of each distribution, and Figure S4b shows their empirical cumulative distribution functions. Figure S4c shows KL divergences of approximate distributions to the true distribution which decay as *ε* → 0. Indeed, these three figures indicate that the approximate distributions converge to the full distribution as *ε* → 0. Figure S4d displays boxplots of the runtime in log-seconds for Algorithm 2 as well as the runtime to compute the full distribution. In this case, we see that the Algorithm 2 becomes slower than computing the full distribution when *ε* ≲ 10^*−*5^.

To assess the effect of sequence lengths on Algorithm 2, we perform the same experiment as above on pairwise aligned VDJ sequences (via the `sequence_alignment` column rather than inferred CDR3 sequences. These length distributions are different by about an order of magnitude. We note that the pairwise aligned VDJ sequences are the default for Algorithm 2 within `sumrep`, although we anticipate users to examine this distribution for CDR3s as well as full V(D)J sequences. We run Algorithm 2 on the same subsampled 10,000 sequences of `p_f1`.

Figure S5a shows a frequency polygon of the same distributions, and Figure S5b shows their empirical cumulative distribution functions. Moreover, Figures S5c and S5d show the KL-divergences to truth and runtimes, respectively. It seems that the KL divergence to the truth may converge more slowly for `sequence_alignment` sequences rather than CDR3s, although the approximate procedure seems to outper-form the full distribution for a slightly larger range of *ε* values (i.e. until *ε* nears 10^*−*6^).

Next we investigate the effect of dataset size on the performance of Algorithm 2. For sample sizes *n ∈* {exp(6),…, exp(10)}, we subsample `p_f1` without replacement to *n* sequences and compute the pairwise distance distribution of CDR3 sequences for the full subsampled dataset as well as those given by tolerances *ε ∈* {0.1,…, 10^*−*5^}. We perform this experiment 5 times for each *n*.

Figures S6a, S6b, and S6c display boxplots of the KL-divergence to truth, runtime, and time efficiency, respectively. There is not an obvious trend in KL divergence to truth for a given tolerance as sample size increases, although the variability is higher for high tolerances. As expected, runtime increases as tolerance decreases, and also increases with the size of the dataset. This is reasonable since each batch iteration of Algorithm 2 must compute the nearest neighbor distance from each sequence in batch *B* to the full repertoire *R*, which certainly increases in time complexity as *R* increases.

Next we look at the efficiency relative to computing the full distribution as defined in Equation 10. Examining the boxplots near *y* = 0 by log(size), we see that for a dataset of size exp(*k*), we would need a tolerance of at least 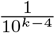. For example, for log(size) = 6, we see that tolerances higher than 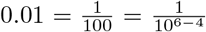 would on average yield an efficiency greater than one. This suggests that, for a dataset with *n* CDR3 sequences, a sensible rule of thumb would be to choose 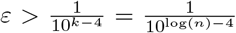. This will of course be more or less appropriate for a given dataset depending on the nature of the repertoire from which it was sampled.

Finally, we perfrom the same experiment but using `sequence_alignment` sequences for the nearest neighbor distance distribution. Figures S7a, S7b, and S7c display boxplots of the KL-divergence to truth, runtime, and time efficiency, respectively. There is evidence of a positive trend of the KL-divergence as sample size increases for *ε* = 0.1, although this trend seems to diminish for each other tolerance. Runtimes increase with given sample size and tolerance, and are generally higher than they are for CDR3 sequences as expected. It turns out that the efficiencies follow the same rule of thumb we derived for the CDR3 sequence situation. In particular, choosing 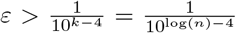 will on average lead to an increase in efficiency with respect to the full distribution for `sequence_alignment` sequences as well as CDR3 sequences. While this may depend on the dataset in question, we recommend this as a good point of reference for general use.

The user should use these results as well as problem-specific considerations when deciding whether or not to use Algorithm 2 instead of computing the full distribution, and if so, which tolerance to use. By default, `sumrep` retrieves the approximate rather than full nearest neighbor distribution, and uses *ε* = 10^*−*4^ unless otherwise modified.

## Appendix C Multinomial lasso path plots

Figure S8 displays the lasso path plots which illustrate the coefficient values of each response vector utilized in Algorithm 3. Figure S8a shows paths for `IGoR` annotations of six TRB datasets from [6], and Figure S8b shows paths for `partis` annotations for six IGH datasets from [18].

## Appendix D Model validation analysis workflows

Figure S9a illustrates the `IGoR` model validation workflow. We employ `IgBLAST` to obtain CDR3 sequences for the observed sequences, which `IGoR` only outputs for generated sequences. Moreover, because we fit `IGoR` models on predominantly productive TRB sequences, we consider only `IGoR`-generated sequences whose V and J segments are in-frame.

Figure S9b illustrates the `partis` model validation workflow as described in the Methods section. We first run `partis partition` on each fasta file of IGH sequence reads to obtain annotations for each sequence, as well as a directory containing model parameters for inference and simulation. We can then run `partis simulate` with these model parameters as input to generate a synthetic datset of IGH annotations. We subsample both the experimental and simulated annotation datasets to unique clones. Then, we compare each IGH-relevant summary for the two resultant annotations datasets, yielding a divergence value for each summary.

## Appendix E Comparison of summary scores using IgBLAST annotations

Recall that for the standard `partis` model validation procedure, `partis` is used for both inference as well as simulation. Here we examine the influence of using the same tool for inference and simulation by using `IgBLAST` for inference, and comparing the annotations dataset output from `IgBLAST` to the corresponding simulations from `partis`. The workflow for this procedure is displayed in Figure S10, which is essentially the diagram in Figure S9b with an additional path describing the `IgBLAST`/Change-O pipeline. Change-O was used to parse the `IgBLAST` output, as well as `partition` the sequences into inferred clonal families [13].

Figure S11a shows the LRAD-data scores by summary when using `IgBLAST` for annotation and `partis` for simulation. Figure S11b shows the difference of each score in Figure 9a and each score in Figure S11a. Frequency polygons of summary distributions of three pairs of `IgBLAST`-annotated and `partis`-simulated datasets are shown in Figure S12. The plots show a high level of agreement for most summaries, with all but six of them differing by less than one units, and a strong majority of them close to zero. Where differences arise, this is likely the result of differences in how `partis` and `IgBLAST` perform annotations. For example, we see that the insertion length distributions highly disagree in scores. This is at least partially attributable to the star-tree assumption on which `partis` operates, which is prone to overestimate insertion lengths in an effort to better estimate the ultimate naive sequence. Indeed, examining the VD insertion length distribution shows that `IgBLAST` tends to assign a similar distribution to each dataset, whereas `partis` leads to more variable distributions with right skew due to the star-tree assumption. Moreover, if `IgBLAST` tends to assign a similar insertion length distribution to every dataset, then this will make it difficult for a simulator designed to match particular insertion lengths distributions to behave more like the `IgBLAST` distributions. Thus, inherent differences in annotation tools will certainly lead to differences in summary scores, regardless of how accurate either tool is. Hence, it is important to understand that a given annotations-based summary should be considered in the context of the tool which provided annotations, and not as a ground-truth summary of the actual gene usage/indel statistics.

---

**Figure S1:**
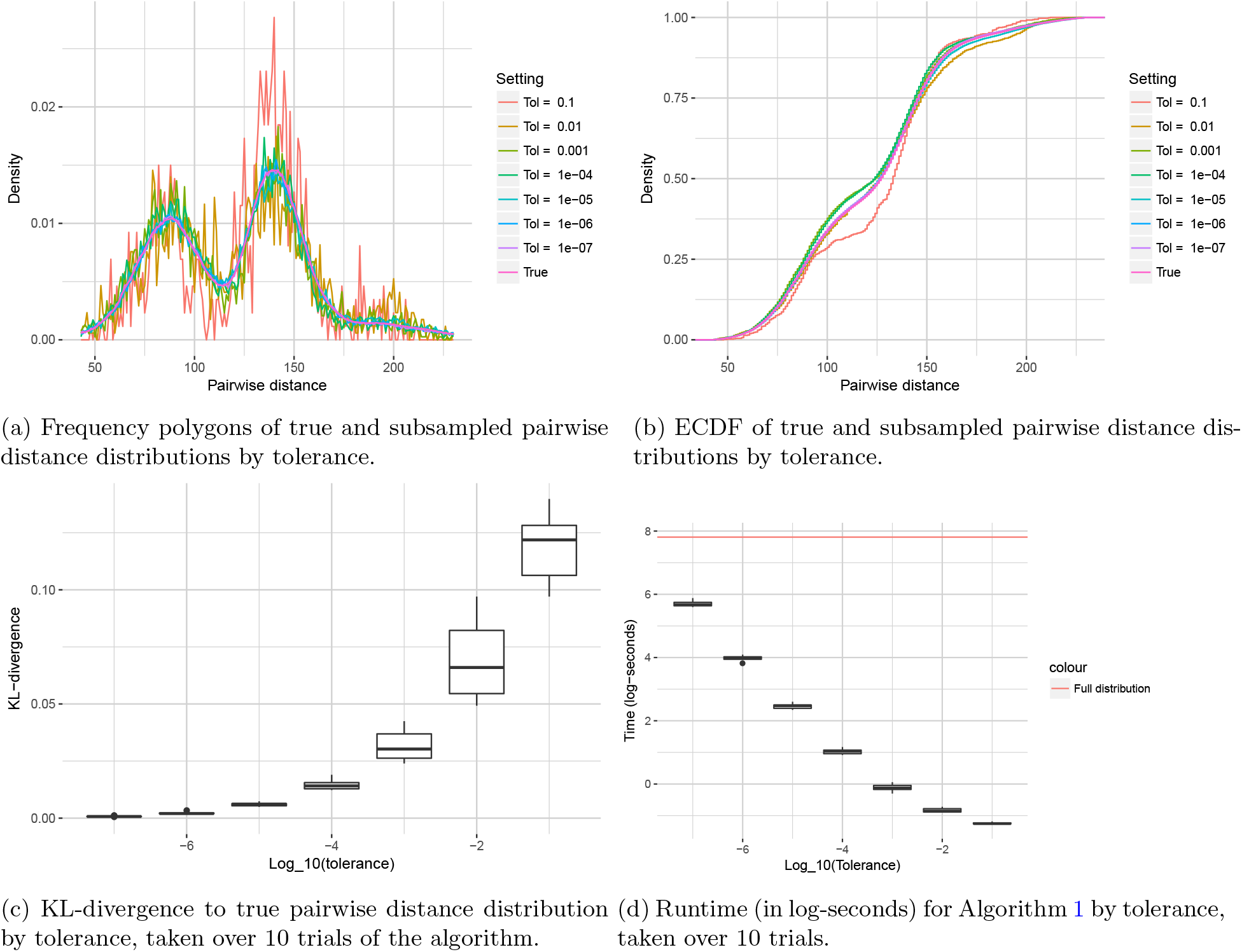
Performance of Algorithm 1 by tolerance applied to the pairwise distance distribution.

**Figure S2:**
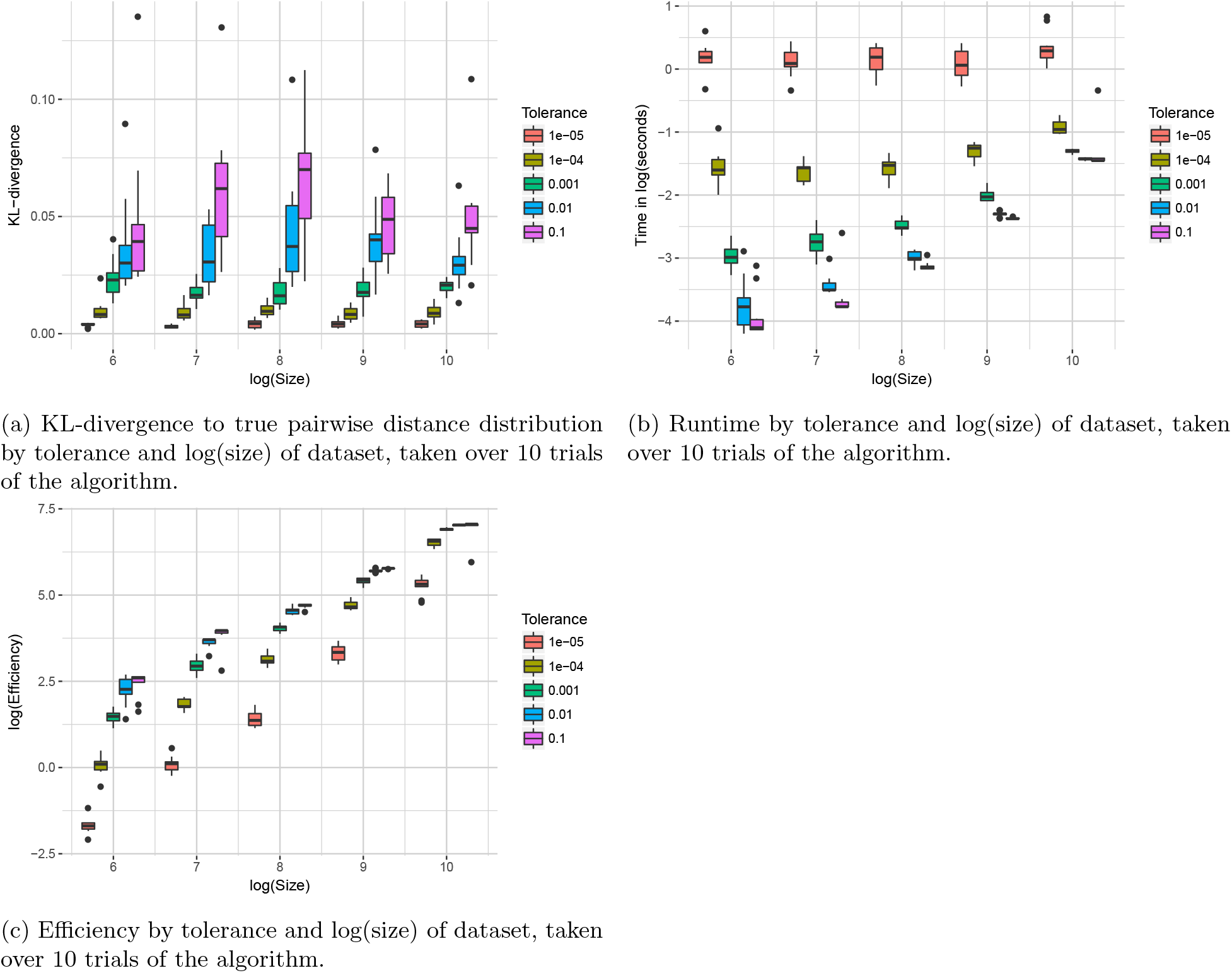
Performance of Algorithm 1 by sample size and tolerance applied to the pairwise distance distribution.

**Figure S3:**
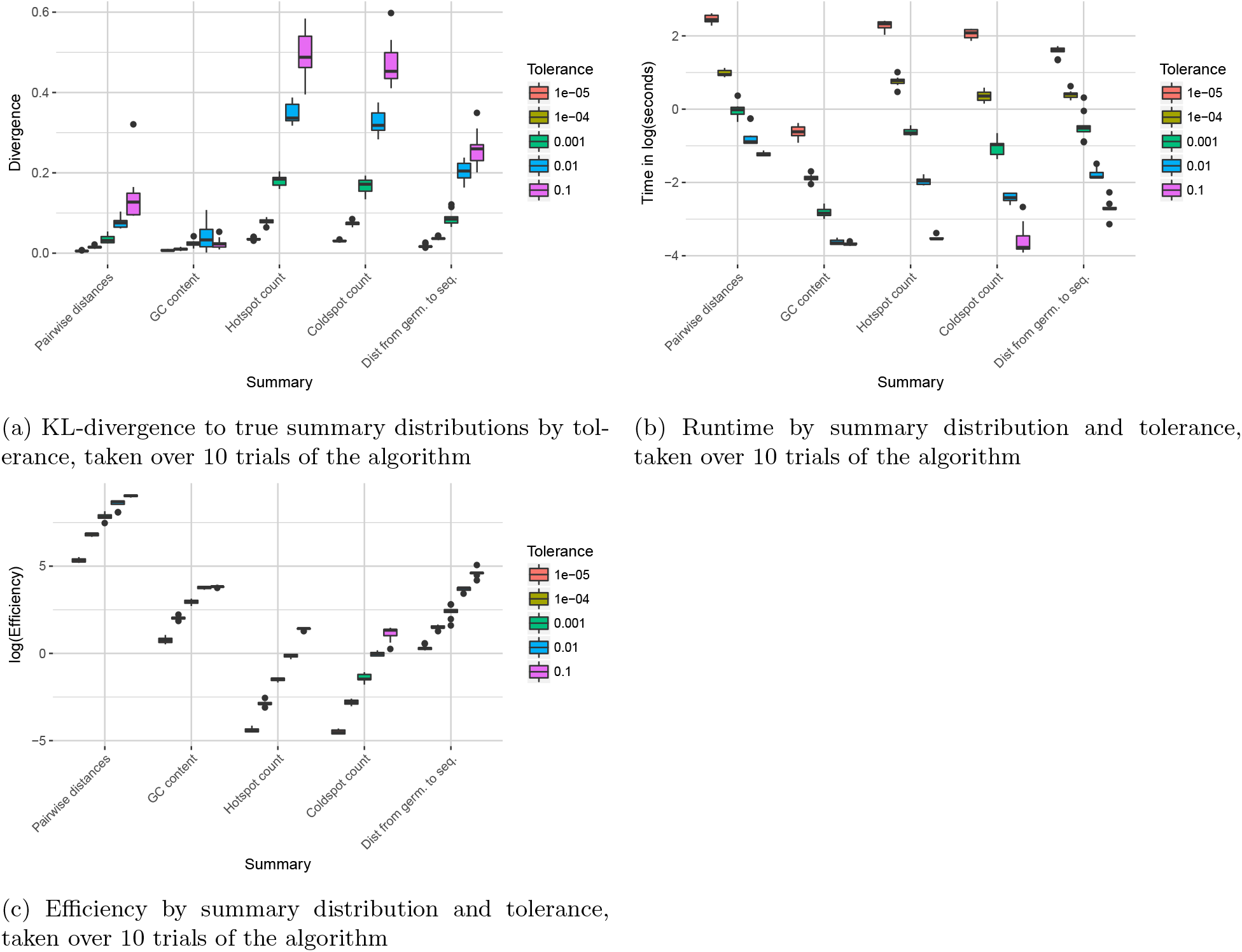
Performance of Algorithm 1 by summary statistic and tolerance applied to the pairwise distance distribution.

**Figure S4:**
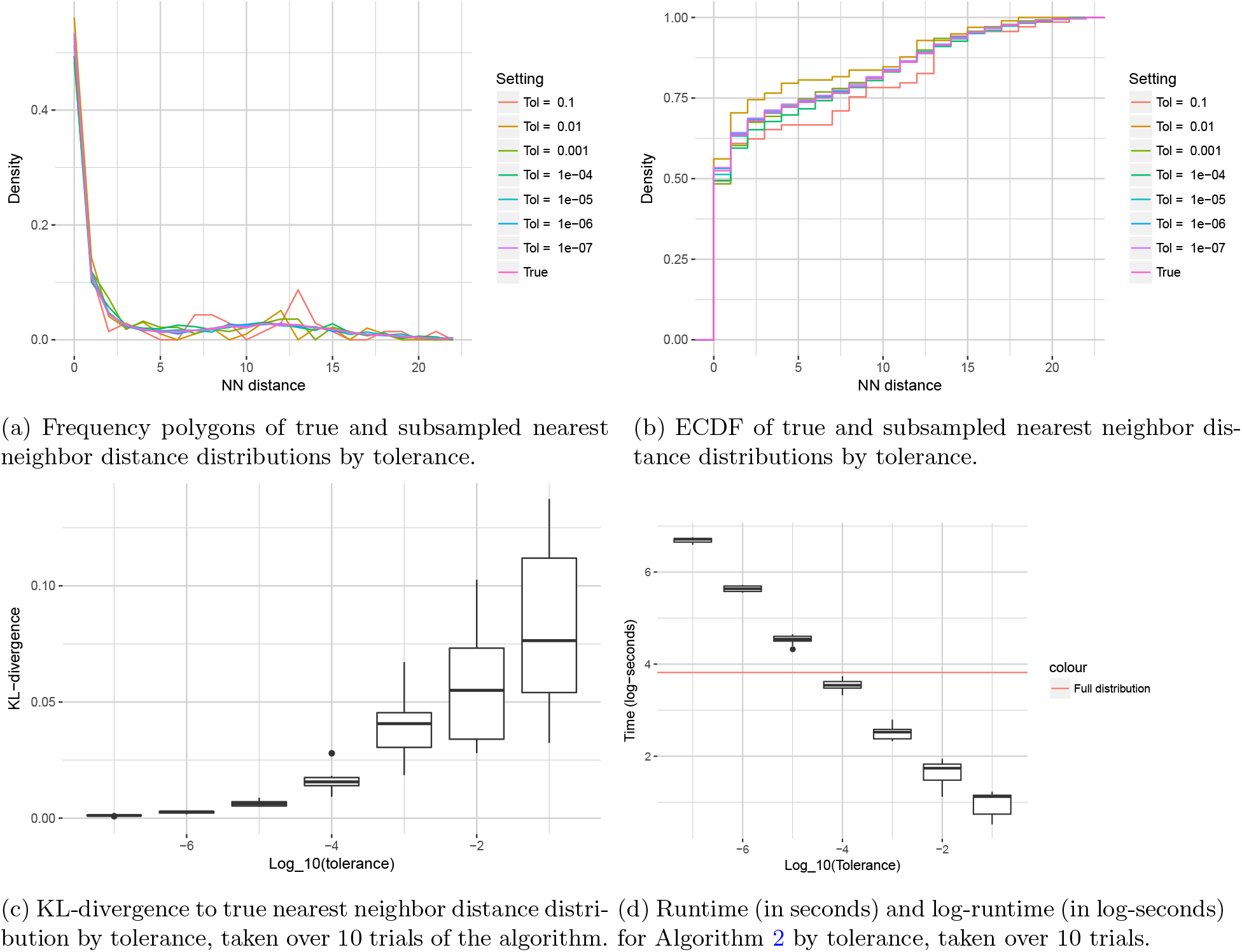
Performance of Algorithm 2 by tolerance applied to the nearest neighbor distribution of CDR3nt sequences.

**Figure S5:**
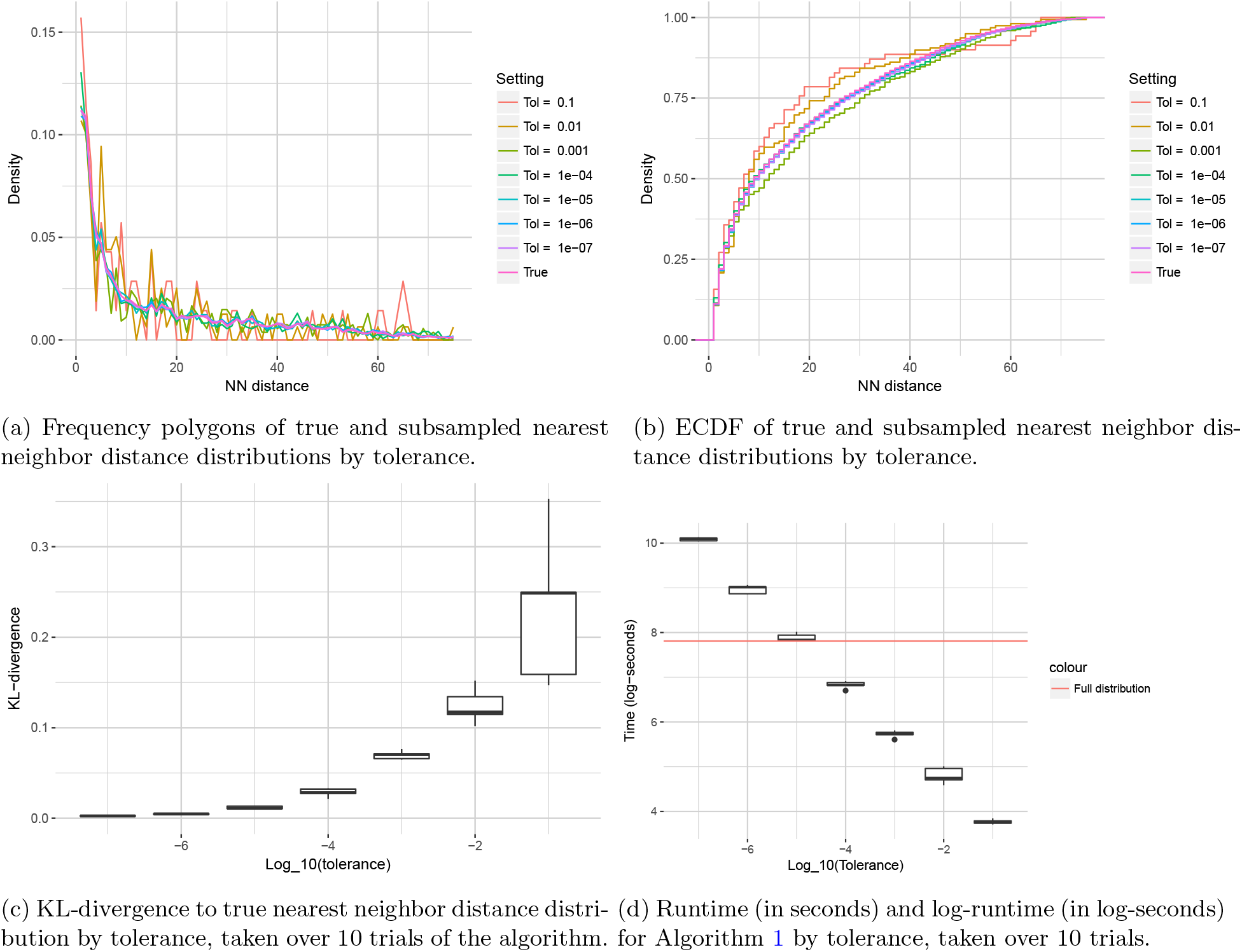
Performance of Algorithm 2 by tolerance applied to the nearest neighbor distribution of pairwise-aligned VDJ sequences.

**Figure S6:**
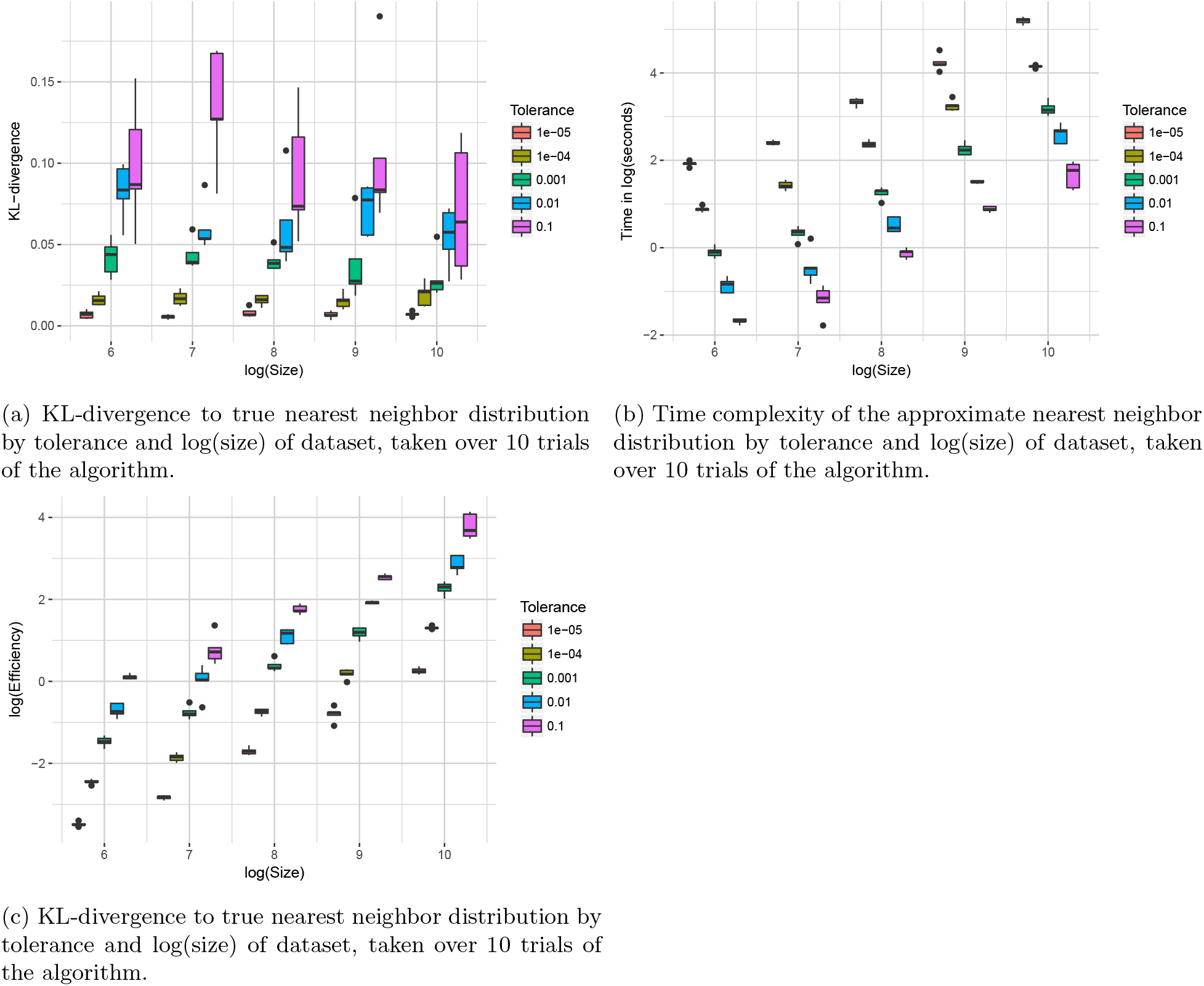
Performance of Algorithm 2 by sample size and tolerance applied to the nearest neighbor distribution of CDR3nt sequences.

**Figure S7:**
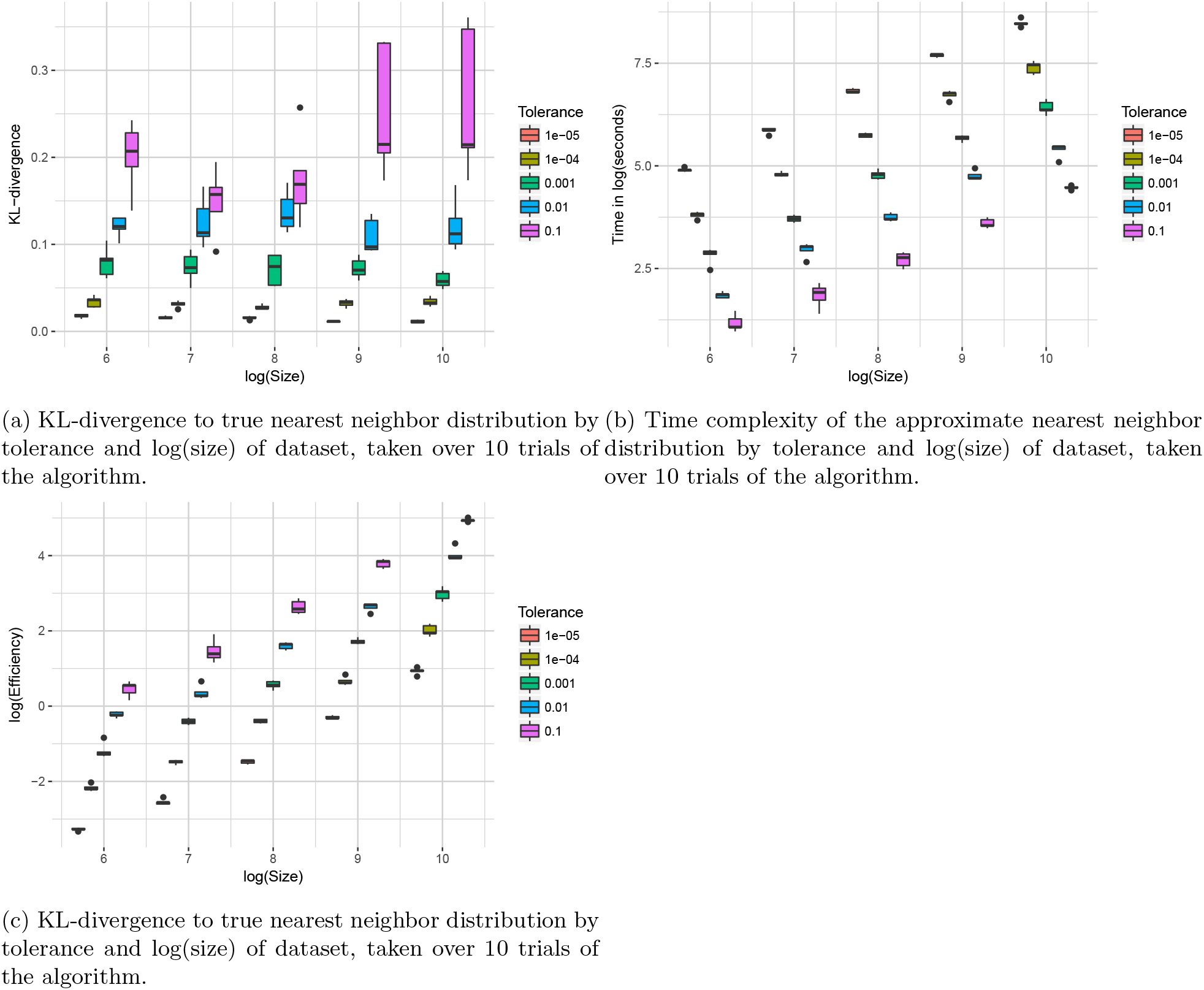
Performance of Algorithm 2 by sample size and tolerance applied to the nearest neighbor distribution of pairwise-aligned VDJ sequences.

**Figure S8:**
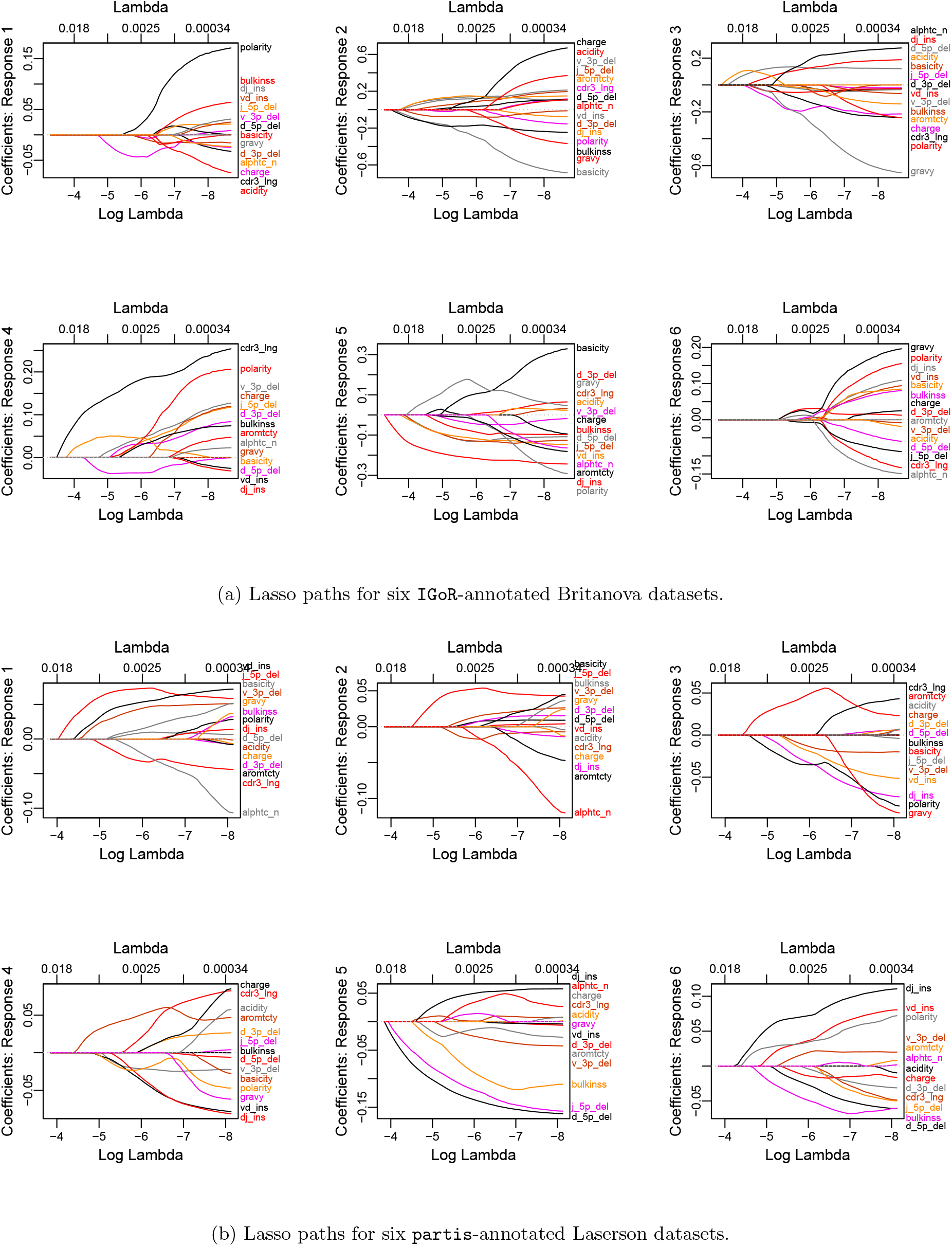
Multinomial lasso paths of summary coefficients by dataset identity.

**Figure S9:**
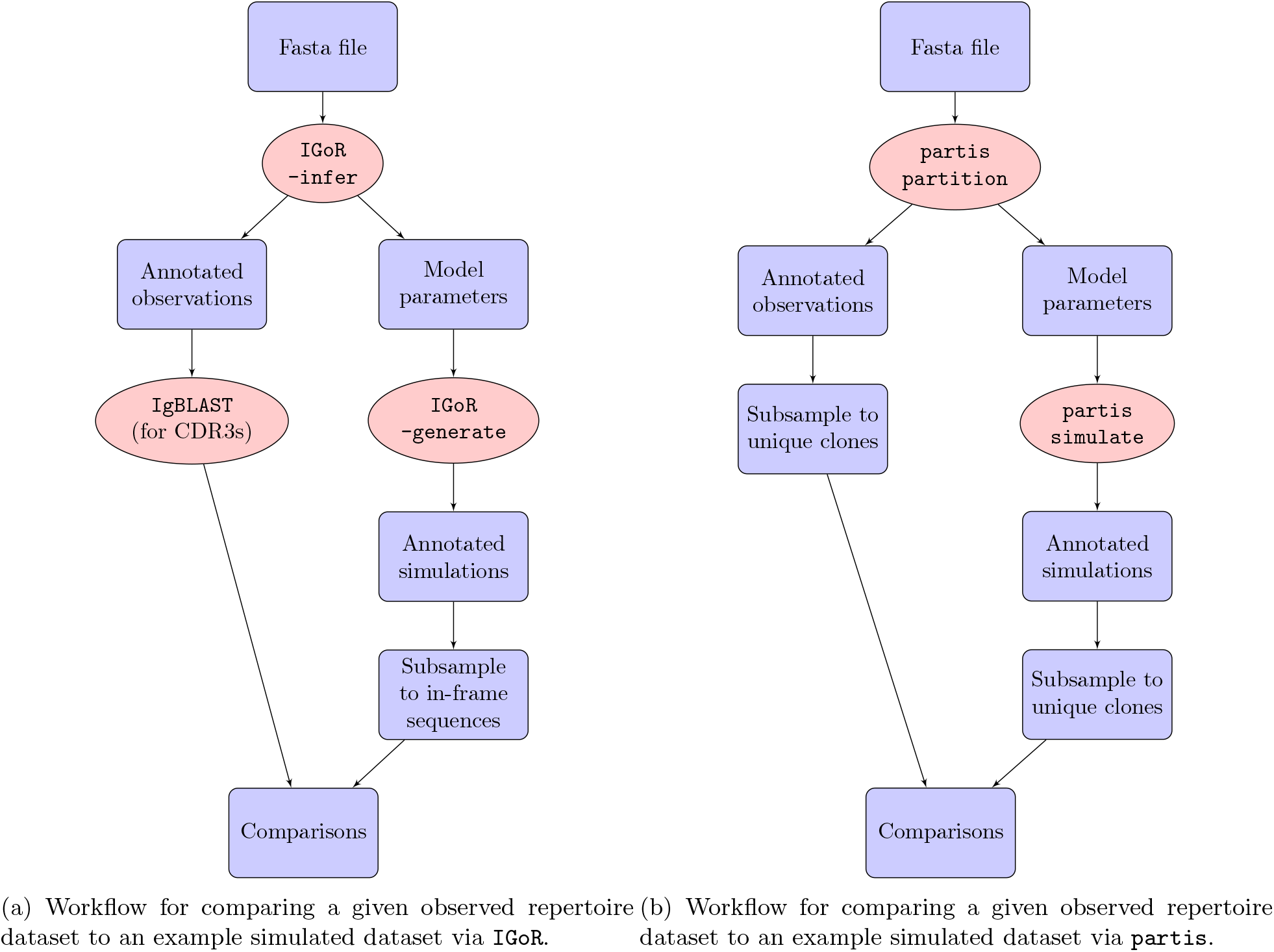
Workflow diagrams for the `IGoR` and `partis` model validation analyses.

**Figure S10:**
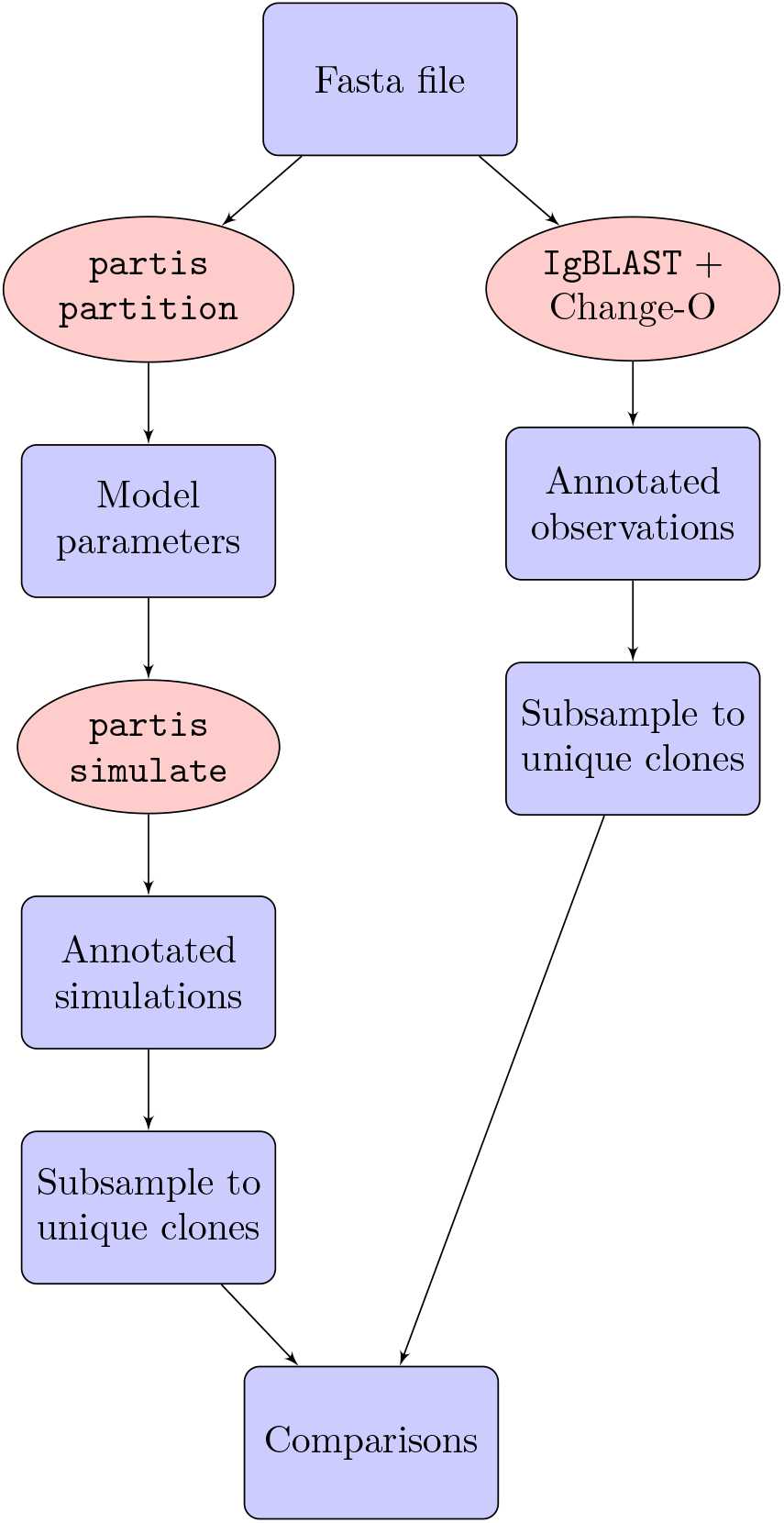
Workflow diagram for `partis` model validation when comparing `partis` and `IgBLAST` annotations to partis simulations

**Figure S11:**
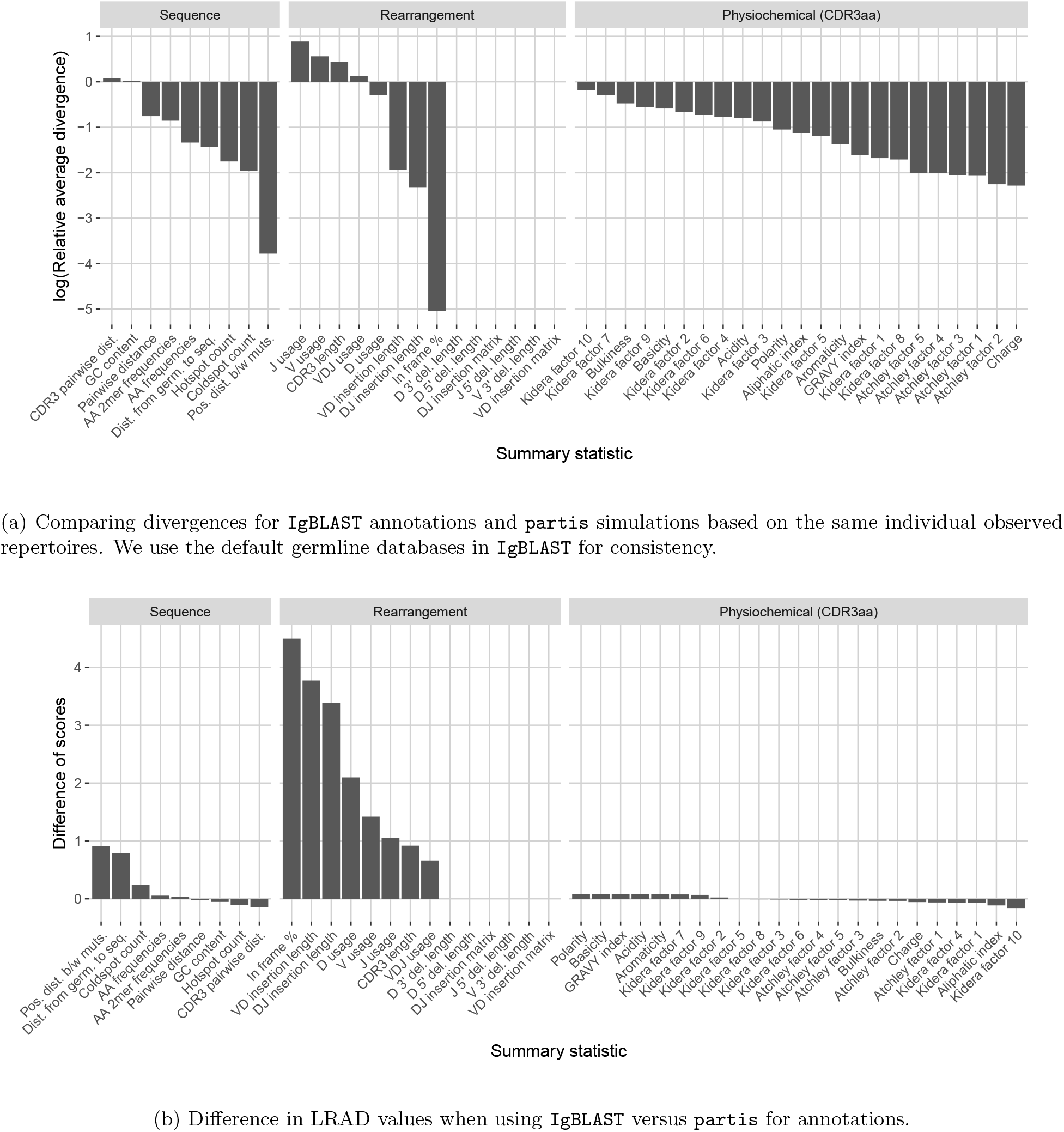
Summary scores for each statistic in the `partis` model validation experiment when comparing `partis` simulations to `IgBLAST` annotations. IMGT IGH germline databases were used during inference for both tools. In both plots, a high score indicates a well-replicated statistic by the simulations with respect to their corresponding experimental repertoires of productive IGH sequences. Summaries without a score are not readily available from AIRR-formatted `IgBLAST` output.

**Figure S12:**
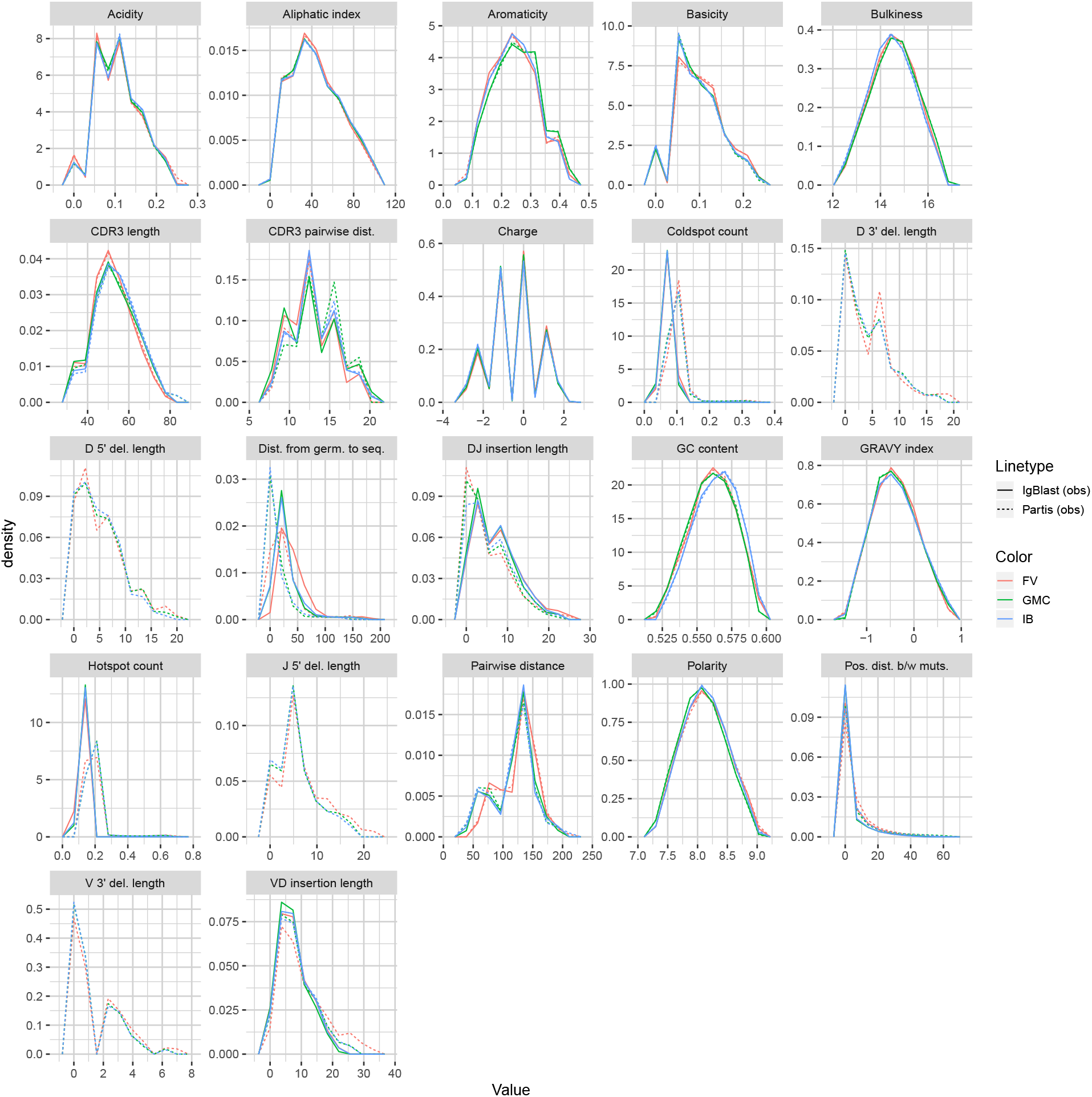
Summary distribution frequency polygons of `partis` versus `IgBLAST` annotations of experimental datasets from three donors at time point −1h.

## References

[1] Oren Avram, Anna Vaisman-Mentesh, Dror Yehezkel, Haim Ashkenazy, Tal Pupko, and Yariv Wine. Asap – a webserver for immunoglobulin-sequencing analysis pipeline. Front. Immunol., 9:1686, July 2018.

[2] Julia Bischof and Saleh M. Ibrahim. bcRep: R package for comprehensive analysis of B cell receptor repertoire data. PLoS ONE, 11(8):e0161569, August 2016.

[3] Carl Boettiger. An introduction to docker for reproducible research. SIGOPS Oper. Syst. Rev., 49(1):71–79, January 2015.

[4] CR Bolen, F Rubelt, JA Vander Heiden, and MM Davis. The repertoire dissimilarity index as a method to compare lymphocyte receptor repertoires. BMC Bioinformatics, 18(1):155, March 2017.

[5] SD Boyd et al. Individual variation in the germline Ig gene repertoire inferred from variable region gene rearrangements. J Immunol, 184(12):6986–92, June 2010.

[6] O V Britanova et al. Dynamics of individual T cell repertoires: from cord blood to centenarians. J. Immunol., 196(12):5005–5013, June 2016.

[7] MM Corcoran, GE Phad, Bernat N Vázquez, C Stahl-Henning, N Sumida, MA Persson, M Martin, and Hedestam GB Karlsson. Production of individualized V gene databases reveals high levels of immunoglobulin genetic diversity. Nat Commun, 7:13642, December 2016.

[8] Marc Duez, Mathieu Giraud, Ryan Herbert, Tatiana Rocher, Mikael Salson, and Florian Thonier. Vidjil: A web platform for analysis of high-throughput repertoire sequencing. PLoS One, 11:e0166126, November 2016.

[9] Yuval Elhanati, Anand Murugan, Curtis G Callan, Jr, Thierry Mora, and Aleksandra M Walczak. Quantifying selection in immune receptor repertoires. Proc. Natl. Acad. Sci. U. S. A., June 2014.

[10] William J. J. Finlay and Juan C. Almagro. Natural and man-made V-gene repertoires for antibody discovery. Front. Immunol., 3:342, 2012.

[11] D Gadala-Maria, G Yaari, M Uduman, and SH Kleinstein. Automated analysis of high-throughput B-cell sequencing data reveals a high frequency of novel immunoglobulin V gene segment alleles. Proc Natl Acad Sci U S A, 112(8):E862–70, February 2015.

[12] Namita T. Gupta, Kristofor D. Adams, Adrian W. Briggs, Sonia C. Timberlake, Francois Vigneault, and Steven H. Kleinstein. Hierarchical clustering can identify b cell clones with high confidence in ig repertoire sequencing data. The Journal of Immunology, 198(6):2489–2499, 2017.

[13] Namita T. Gupta, Jason A. Vander Heiden, Mohamed Uduman, Daniel Gadala-Maria, Gur Yaari, and Steven H. Kleinstein. Change-O: a toolkit for analyzing large-scale B cell immunoglobulin repertoire sequencing data. Bioinformatics, 31(20):3356–3358, October 2015.

[14] James M Heather, Mattia Cinelli, Benny Chain, Katharine Best, Yuxin Sun, John Shawe-Taylor, Eric Shifrut, Shlomit Reich-Zeliger, and Nir Friedman. Feature selection using a one dimensional naive Bayes classifier increases the accuracy of support vector machine classification of CDR3 repertoires. Bioinformatics, 33(7):951–955, 01 2017.

[15] D Hou et al. Immune repertoire diversity correlated with mortality in avian influenza A (H7N9) virus infected patients. Sci Rep, 6:33843, September 2016.

[16] Hanna IJspeert, Pauline A van Schouwenburg, David van Zessen, Ingrid Pico-Knijnenburg, Andrew P Stubbs, and Mirjam van der Burg. Antigen receptor galaxy: A user-friendly, web-based tool for analysis and visualization of t and b cell receptor repertoire data. J. Immunol., 198:4156–4165, May 2017.

[17] Kevin Larimore, Michael W McCormick, Harlan S Robins, and Philip D Greenberg. Shaping of human germline IgH repertoires revealed by deep sequencing. J. Immunol., 189(6):3221–3230, August 2012.

[18] Uri Laserson, Francois Vigneault, Daniel Gadala-Maria, Gur Yaari, Mohamed Uduman, Jason A. Vander Heiden, William Kelton, Sang Taek Jung, Yi Liu, Jonathan Laserson, Raj Chari, Je-Hyuk Lee, Ido Bachelet, Brendan Hickey, Erez Lieberman-Aiden, Bozena Hanczaruk, Birgitte B. Simen, Michael Egholm, Daphne Koller, George Georgiou, Steven H. Kleinstein, and George M. Church. High-resolution antibody dynamics of vaccine-induced immune responses. PNAS, 111(13):4928–4933, April 2014.

[19] Asaf Madi, Benny Chain, Eric Shifrut, Hilah Gal, John Shawe-Taylor, Katharine Best, Mattia Cinelli, Niclas Thomas, Nir Friedman, and Shlomit Reich-Zeliger. Tracking global changes induced in the CD4 T-cell receptor repertoire by immunization with a complex antigen using short stretches of CDR3 protein sequence. Bioinformatics, 30(22):3181–3188, 08 2014.

[20] Quentin Marcou, Thierry Mora, and Aleksandra M. Walczak. High-throughput immune repertoire analysis with IGoR. Nat. Commun, 9(561), 2018.

[21] V Martin, YC Bryan Wu, D Kipling, and D Dunn-Walters. Ageing of the B-cell repertoire. Philos Trans R Soc Lond B Biol Sci, 370(1676), September 2015.

[22] Lisa McFerrin. HDMD: Statistical Analysis Tools for High Dimension Molecular Data DMD, 2013. R package version 1.2.

[23] P Miqueu, M Guillet, N Degauque, JC Doré, JP Soulillou, and S Brouard. Statistical analysis of CDR3 length distributions for the assessment of T and B cell repertoire biases. Mol Immunol, 44(6):1057–1064, February 2007.

[24] Arnau Mir, Francesc Rossello, and Lucia Rotger. CollessLike: Distribution and Percentile of Sackin, Cophenetic and Colless-Like Balance Indices of Phylogenetic Trees, 2018. R package version 1.0.

[25] Vadim I Nazarov, Mikhail V Pogorelyy, Ekaterina A Komech, Ivan V Zvyagin, Dmitry A Bolotin, Mikhail Shugay, Dmitry M Chudakov, Yury B Lebedev, and Ilgar Z Mamedov. tcr: an R package for T cell receptor repertoire advanced data analysis. BMC Bioinformatics, 16(1):175, May 2015.

[26] Jared Ostmeyer, Scott Christley, William H. Rounds, Inimary Toby, Benjamin M. Greenberg, Nancy L. Monson, and Lindsay G. Cowell. Statistical classifiers for diagnosing disease from immune repertoires: a case study using multiple sclerosis. BMC Bioinformatics, 18(1):401, Sep 2017.

[27] H. Pagás, P. Aboyoun, R. Gentleman, and S. DebRoy. Biostrings: String objects representing biological sequences, and matching algorithms, 2017. R package version 2.44.2.

[28] E. Paradis, J. Claude, and K. Strimmer. APE: analyses of phylogenetics and evolution in R lanugage. Bioinformatics, 20(2):289–290, January 2004.

[29] Mikhail V. Pogorelyy, Anastasia A. Minervina, Maximilian Puelma Touzel, Anastasiia L. Sycheva, Ekaterina A. Komech, Elena I. Kovalenko, Galina G. Karganova, Evgeniy S. Egorov, Alexander Yu. Komkov, Dmitriy M. Chudakov, Ilgar Z. Mamedov, Thierry Mora, Aleksandra M. Walczak, and Yuri B. Lebedev. Precise tracking of vaccine-responding t cell clones reveals convergent and personalized response in identical twins. Proceedings of the National Academy of Sciences, 115(50):12704–12709, 2018.

[30] Duncan K. Ralph and Frederick A. Matsen IV. Consistency of VDJ rearrangement and substitution parameters enables accurate B cell receptor sequence annotation. PLOS Comput. Biol., 12(1), January 2016.

[31] Duncan K. Ralph and Frederick A. Matsen IV. Likelihood-based inference of B cell clonal families. PLOS Comput. Biol., 12(10), October 2016.

[32] Florian Rubelt, Christopher R. Bolen, Helen M. McGuire, Jason A. Vander Heiden, Daniel Gadala-Maria, Mikhail Levin, Ghia M. Euskirchen, Murad R. Mamedov, Gary E. Swan, Cornelia L. Dekker, Lindsay G. Cowell, Steven H. Kleinstein, and Mark M. Davis. Individual heritable differences result in unique cell lymphocyte receptor repertoires of naïve and antigen-experienced cells. Nature Communications, 7:11112 EP –, 03 2016.

[33] Susanne Schaller, Johannes Weinberger, Raul Jimenez-Heredia, Martin Danzer, Rainer Oberbauer, Christian Gabriel, and Stephan M Winkler. Immunexplorer (imex): a software framework for diversity and clonality analyses of immunoglobulins and t cell receptors on the basis of imgt/highv-quest preprocessed ngs data. PLoS One, 16:252, August 2015.

[34] Mikhail Shugay, Dmitriy V Bagaev, Maria A Turchaninova, Dmitriy A Bolotin, Olga V Britanova, Ekaterina V Putintseva, Mikhail V Pogorelyy, Vadim I Nazarov, Ivan V Zvyagin, Vitalina I Kirgizova, Kirill I Kirgizov, Elena V Skorobogatova, and Dmitriy M Chudakov. VDJtools: Unifying post-analysis of T cell receptor repertoires. PLOS Comput. Biol., 11(11):e1004503, November 2015.

[35] Mark P.J. van der Loo. The stringdist package for approximate string matching. The R Journal, 6(1):111–122, June 2014.

[36] Jason Anthony Vander Heiden, Susanna Marquez, Nishanth Marthandan, Syed Ahmad Chan Bukhari, Christian E Busse, Brian Corrie, Uri Hershberg, Steven H Kleinstein, Frederick A Matsen, Iv, Duncan K Ralph, Aaron M Rosenfeld, Chaim A Schramm, AIRR Community, Scott Christley, and Uri Laserson. AIRR community standardized representations for annotated immune repertoires. Front. Immunol., 9:2206, September 2018.

[37] Gur Yaari, Mohamed Uduman, and Steven H Kleinstein. Quantifying selection in high-throughput immunoglobulin sequencing data sets. Nucleic Acids Res., 40(17):e134, May 2012.

[38] Jian Ye, Ning Ma, Thomas L. Madden, and James M. Ostell. IgBLAST: an immunoglobulin variable domain sequence analysis tool. Nucleic Acids Res., 41(W1):W34–W40, July 2013.

[39] Ryo Yokota, Yuki Kaminaga, and Tetsuya J. Kobayashi. Quantification of inter-sample differences in T-cell receptor repertoires using sequence-based information. Frontiers in Immunology, 8:1500, 2017.

